# Dantrolene inhibition of ryanodine receptor 1 carrying the severe malignant hyperthermia mutation Y522S visualized by cryo-EM

**DOI:** 10.1101/2024.10.21.619310

**Authors:** Kavita A. Iyer, Takuya Kobayashi, Takashi Murayama, Montserrat Samsó

## Abstract

Mutations in the skeletal isoform of the ryanodine receptor (RyR1) pose grave risks during anesthesia or treatment with succinylcholine muscle relaxants. These can trigger a potentially lethal malignant hyperthermia (MH) episode via intracellular calcium increase mainly from RyR1 channel leakage. Dantrolene is the primary antidote to prevent death. The main target of dantrolene is RyR1, however little is known about the mechanism of inhibition. Cryo-EM of dantrolene bound to the severe MH Y522S RyR1 mutant in the closed and open states at 2.5-3.3 Å resolution revealed that the drug binds to the channel’s cytoplasmic assembly, far from the ion gate, interacting with residues W882, W996 and R1000 in the P1 domain. The finding was validated by Ca^2+^ imaging and [^3^H]ryanodine binding in WT and alanine mutants. Dantrolene reduced channel opening probability by restricting the central activation module, “cooling down” the primed conformation caused by the mutation. These findings shed light on dantrolene’s mechanism against MH and suggest avenues for developing more effective therapeutics.

## INTRODUCTION

Missense mutations in the skeletal muscle ryanodine receptor (RyR1), large intracellular calcium channels that play a well-characterized role in excitation-contraction coupling, cause disorders such as malignant hyperthermia (MH) and central core disease (CCD) (*1-3*). CCD is identified as a muscular disorder marked by central cores in skeletal muscle and reduced force (*4*). MH episodes are caused by exposure to volatile halogenated anesthetics or succinylcholine. These episodes are characterized by heightened muscle metabolism, increased temperature, rhabdomyolysis, and if untreated, can be fatal (*5*). A hallmark of MH and most CCD muscle fibers is an elevated cytosolic resting free Ca^2+^ concentration (*6*) caused by gain-of-function leaky RyR1 channels (*7, 8*). Indeed, RyR1 triggers the Ca^2+^ signals for muscle cell contraction, but also orchestrates Ca^2+^ homeostasis at rest (*9*). The Ca^2+^ dysregulation resulting from RyR1 mutations elevates the resting cytosolic Ca^2+^ concentration set point, decreases endoplasmic reticulum (ER) Ca^2+^ content, and indirectly increases sarcolemmal Ca^2+^ permeability, altogether resulting in a chronic metabolic burden. This renders the muscle cells incapable of effectively responding to triggering conditions which further exacerbate the elevated intracellular Ca^2+^ in MH-susceptible muscles (*10-13*).

Dantrolene, originally developed as a muscle relaxant (*14*), was tested for the treatment of MH crises first in swine (*15-17*) and later used in humans, where prompt administration of dantrolene upon the onset of an MH episode was shown to be effective in saving patients (*18*). Despite drawbacks such as poor water solubility, dantrolene continues to remain the only clinically approved drug to treat MH episodes for the past 5 decades (*19*). Subsequently, since the 1980s availability of dantrolene in operating rooms is mandatory (*20*). In 2014, the U.S. Food and Drug Administration (FDA) approved Ryanodex (Eagle Pharmaceuticals, Inc.), a new formulation of dantrolene requiring substantially lesser volumes and faster reconstitution (*21*), indicating continued interest in dantrolene and a lack of newer drugs. Dantrolene is also a muscle relaxant indicated for muscle spasticity. Its advantage is that unlike other hydantoin derivatives, dantrolene does not act through the central nervous system and causes less sedation (*22*).

The main target of dantrolene is RyR1, where dantrolene acts as a selective inhibitor (*23, 24*). An additional role in attenuating extracellular Ca^2+^ entry was also proposed (*25*). Despite its known efficacy, the binding site and mechanism of inhibition of RyR1 by dantrolene have remained inconclusive, with different binding sites proposed (*26, 27*). Structural elucidation of dantrolene’s site of interaction with the RyR1 and identifying any ensuing conformational alterations can yield tremendous insight on its mechanism of action and aid in the development of more effective inhibitors.

Recent advances in cryo-electron microscopy (cryo-EM) have allowed a better understanding of the molecular mechanisms of single-point MH-causing mutations such as R164C (*28*), Y523S (*29*), and R615C (*30*). The common trend observed in all these studies is that mutant channels in closed-state conditions adopt a conformation half-way to the open state, thus changing the conformational landscape of RyR1, such that it requires less energy to open. Quantitative analysis of the effect of the severe Y523S RyR1 MH/CCD mutation pinpointed the molecular mechanism and showed the large-scale conformational changes which explain the increased sensitization of the channels to activation (*29*). This mutation also affected the open conformation, which acquired augmented open-state characteristics.

Here we unravel the mechanism of inhibition of dantrolene by determining the structure of the well-characterized Y523S RyR1 mutant in the presence of dantrolene under closed- and open-state conditions (i.e., absence and presence of activating calcium, respectively, while all other conditions remain the same). Systematic and quantitative comparisons with the cryo-EM structures of RyR1 Y523S in the absence of dantrolene, previously determined by us (PDB IDs: <u>7T64</u> and <u>7T65</u>), reveal the binding site of dantrolene on RyR1 and shed light on the mechanism by which dantrolene achieves inhibition of the defective channel, consisting of partial reversal of the conformational anomaly created by the mutation.

## RESULTS

We sought to determine the reversing effects of dantrolene on the structure of a previously characterized gain-of-function (leaky) mutant RyR1, Y523S (rabbit sequence, equivalent to Y522S in human). Our previous work revealed that these mutant channels were more prone to activation by the functional tests of ryanodine binding and live cell Ca^2+^ imaging (*31*), and displayed a noticeable altered conformation when examined by cryo-EM. In the closed state, Y523S RyR1 adopted a closed-primed conformation, while the open-state conformation was amplified (*29*).

Here we determined the structure of RyR1 Y523S-FKBP12.6-dantrolene complex in the closed state without Ca^2+^ (2 mM EGTA) and in the open state with Ca^2+^ present (50 μM free Ca^2+^), henceforth referred to as RyR1^YS^-closed-DAN and RyR1^YS^-open-DAN, respectively. As in the case of the RyR1^YS^ mutants alone, both 3D reconstructions were determined in the presence of ATP, a constitutive ligand of RyR1 in vivo. The transmembrane domain of RyR1 was stabilized with nanodiscs. The global resolutions of the cryo-EM structures were 3.3 Å and 3.2 Å (FSC_0.143_) for RyR1^YS^-closed-DAN and RyR1^YS^-open-DAN, respectively, with large parts (∼30 %) of the structures reaching ∼2.5 Å (Figs. S1-S5, Table S1). The channels display the prototypical prism-shaped cytoplasmic assembly (CytA) and stalk-like transmembrane domain (TmD), and well-defined individual domains as described previously (*29, 32*); inset in Fig. S1. ATP was clearly discernable on both structures (Fig. S6A) and the Ca^2+^ binding site was occupied only in the open state (Fig. S6B).

### Cryo-EM reveals that dantrolene binds to the corners of RyR1

A striking feature of the 3D reconstructions upon addition of dantrolene was the closure of the V-shaped clamp of the P1 domain (residues 847-1055) situated at the four corners of the prism (Fig. 1A and S7), accompanied by extra density within the clamp. By virtue of their position far from the structure’s center, the P1 domains suffer most from misalignment and had poor resolution, which was overcome by focused alignment. The focused maps for the P1 region of the closed and open reconstructions, with resolutions of 3.8 and 3.2 Å, respectively, revealed a distinct density that fitted closely the dantrolene molecule (Fig. 1B). Dantrolene nestled in the cavity at the intersection of the 4 α helices interacting with charged (E917 and R1000), polar (N921, N1035, S1038) and hydrophobic residues (I878, W882, Y920, M924, and W996) (Figs. 1C and S7A-B). The phenyl ring of the dantrolene molecule engages in a parallel-displaced π-π stacking interaction with the indole ring of a neighboring tryptophan residue, W996, while the furane group of dantrolene faces the indole ring of W882 (Fig. 1D and Fig. S7A). The oxygen atom of the carbonyl group on the imidazolidine ring is in the vicinity of the positively charged guanidinium moiety of R1000. Of note, it has been well documented that mutation R1000H causes CCD (*33*). Residues S1038 or N1035 can interact with the nitro group on the phenyl ring of dantrolene via a hydrogen bond (Figs. 1C and S7B). The resolution of the density corresponding to dantrolene is such that it can be modeled either with the nitro end or with the imidazolidine ring sticking into the cleft towards M924. However, in the orientation with the imidazolidine ring facing M924, the phenyl ring is no longer able to effectively interact with W882 in a π-π stacking interaction. Minor differences in orientation of the dantrolene molecule were observed between the RyR1^YS^-closed-DAN and RyR1^YS^-open-DAN. In the RyR1^YS^-open-DAN map, the nitro group forms a hydrogen bond with S1038, while the nitrogen atom of the indole ring of W882 forms a cation-π interaction with the imidazoline ring of dantrolene. All these interactions mediated by dantrolene pulled in helix H2 (residues 915-933) and its directly connected helix H1 (residues 865-881), which form the outer arm of the V, by 10 Å (Figs. 1 and S7C). In addition, we observed partial density in the well-documented caffeine site of RyR1, located at the intersection of CD, CTD, U-motif of one subunit and S2-S3 linker of the neighboring subunit (*34*). The partial density appears as an elongated lump sandwiched between tryptophan residues, W5011 of CTD and W4716 of S2-S3 linker in the RyR1^YS^-closed-DAN map (Fig. S8A). Modeling of dantrolene at this secondary site was made difficult since the density disappeared at the threshold used to visualize the contour of the surrounding side chains. Conformational change in this region was negligible, suggesting a potential secondary site of lesser importance. We also examined the region around the DP1 peptide (residues 590-609) known to bind dantrolene as a synthetic peptide (*26*). All density in its vicinity was accounted by the side chains, and while lowering the threshold revealed elongated extra densities facing residues 595 and 596, these are too small to accommodate the dantrolene molecule (Fig. S8B). Thus, it appears that the affinity of dantrolene for the DP1 peptide is substantially lower when the peptide forms part of RyR1. Relative locations of the putative secondary site and DP1 are shown in Fig. S8C.

**Figure 1.**
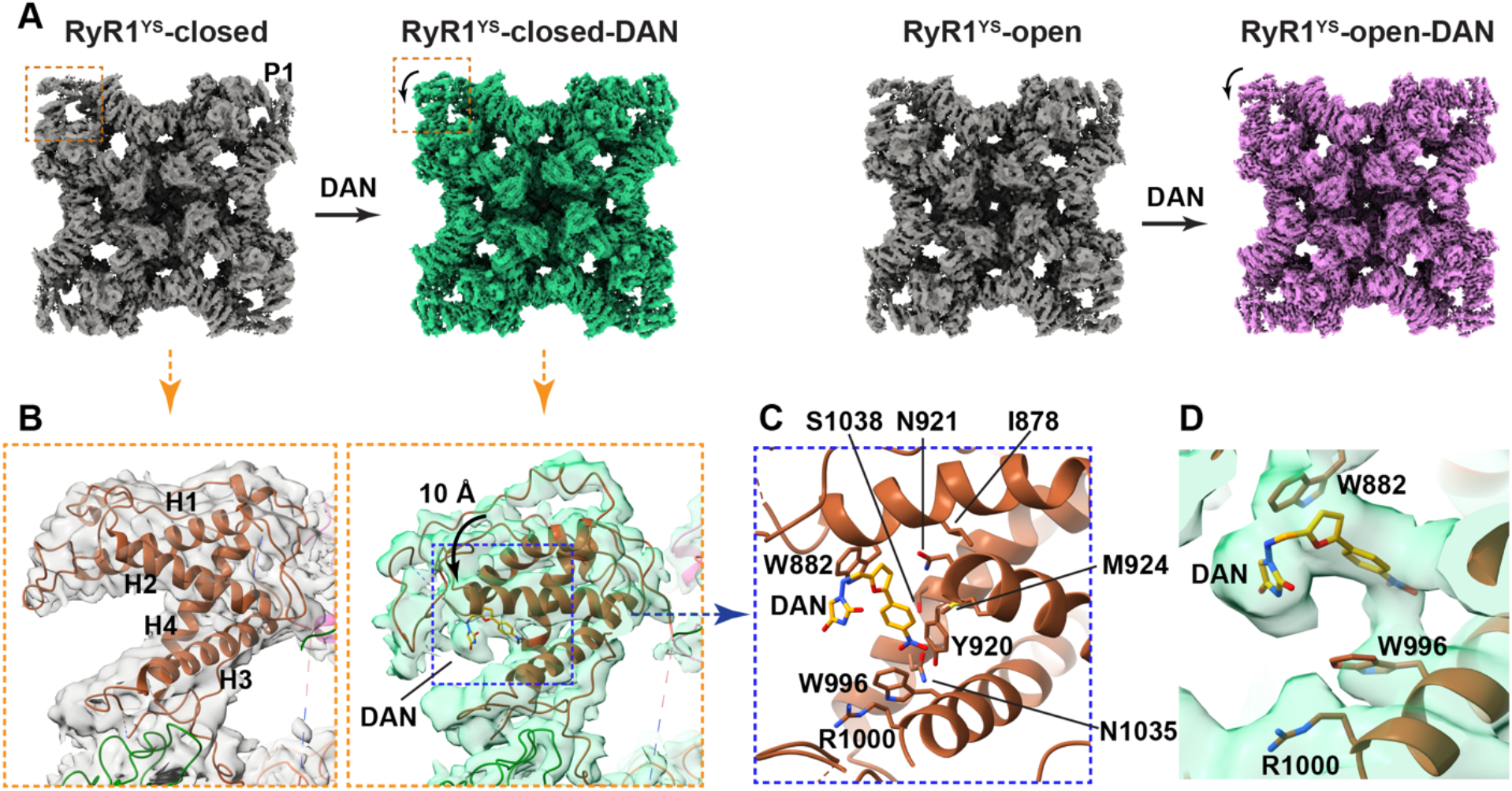
Binding site and effect of dantrolene on the 3D conformation of RyR1^YS^. (**A**) The conformational change of the P1 domain upon addition of dantrolene is shown for the closed state (left; green) and open state (right; magenta) conditions; mutant RyR1s^YS^ under the same conditions are shown in gray for reference. In the presence of dantrolene, the P1 domain gets pulled towards the main body of the channel. (**B**) Enlarged region within the dashed squares in (A). Extra density corresponding to dantrolene (DAN) was observed nestled between the V-shaped crevice of the P1 domain when compared to the cryo-EM density of the mutant channel in the absence of dantrolene. The cryo-EM density is shown in semitransparent color with the dantrolene molecule in orange. The same observations apply to the open state (Fig. S7). (**C**) Detail of the region within the blue square in (B) showing residues in the immediate vicinity of dantrolene; see Fig. S7 for more detail. (**D**) Cryo-EM density showing the disposition of the functionally validated dantrolene interacting-residues.

### Validation of the dantrolene binding site in RyR1

To validate the binding site as well as the orientation of dantrolene, we generated the following single point RyR1 mutants: W882A, W996A and R1000A, and expressed them in HEK293 cells (Fig 2). We used the fluorescent probe R-CEPIA1er (*35*), sensitive to Ca^2+^ concentration in the ER/SR compartment, which provides a direct measure of how dantrolene, by reducing the constitutive leak of Ca^2+^ via RyR1 (*9*), increases ER/SR Ca^2+^ content. Adding dantrolene to HEK293 cells expressing WT RyR1 dose-dependently increased the ER Ca^2+^ as shown earlier (*35*) (Fig. 2A, B). In contrast, adding dantrolene to W882A RyR1-expressing cells did not have any effect (Fig. 2A, B). An intermediate effect was observed for W996A and R1000A RyR1-expressing cells, which required >10-fold higher concentration of dantrolene (Fig. 2A, B). Calculation of IC_50_ values revealed that W996A and R1000A RyR1 were 30- and 10-fold, respectively, less sensitive to dantrolene compared to WT RyR1; i.e., dantrolene was less potent at increasing ER/SR Ca^2+^ content (Fig. 2C). These results reflect the relative importance of each of these residues on the affinity to dantrolene, with W882 being essential, and W996 and R1000 also being critical.

**Figure 2.**
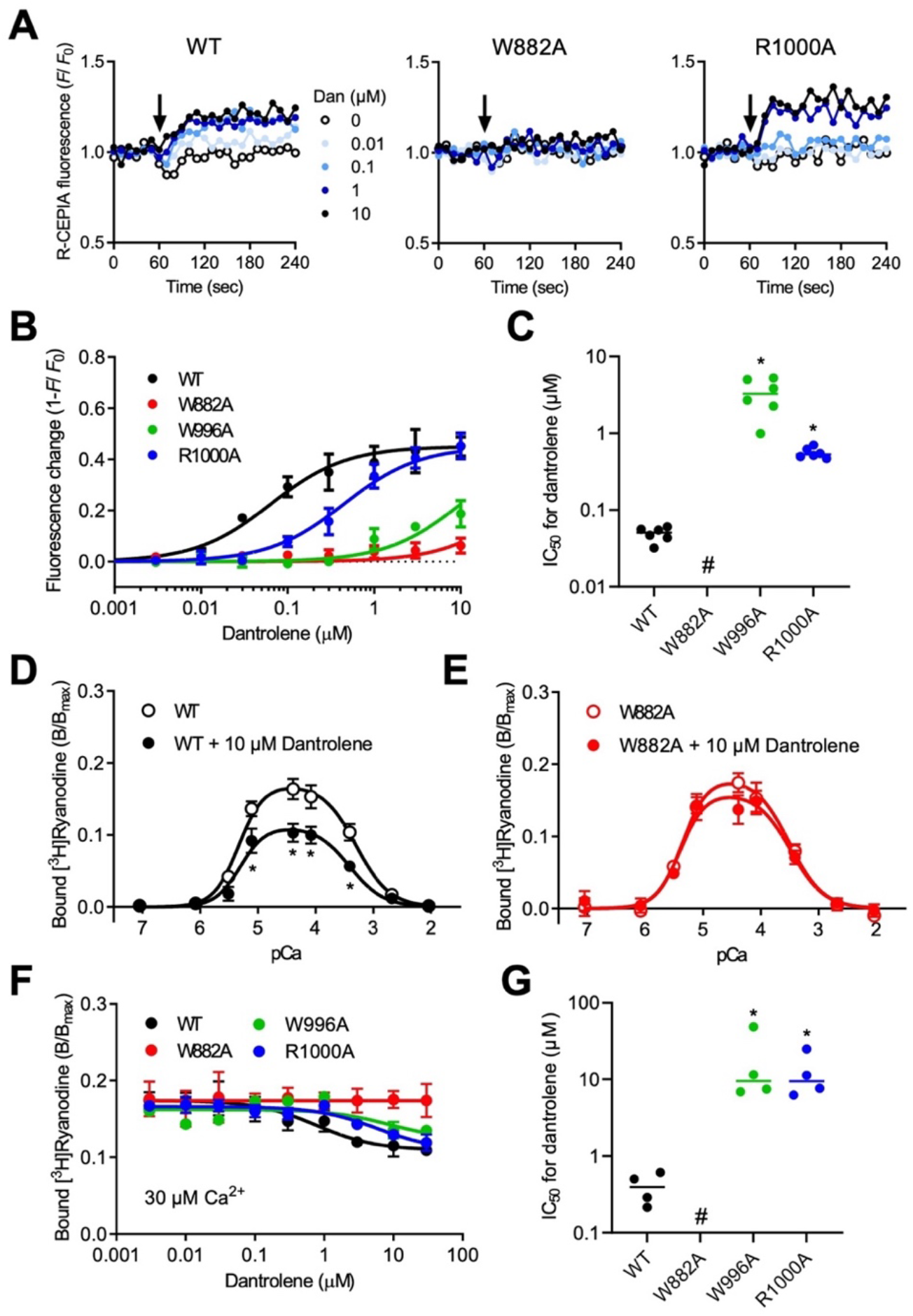
Functional validation of RyR1 channels carrying mutations in the putative dantrolene-binding site. (**A-C)**. Fluorescent measurement of [Ca^2+^]_ER_ in HEK293 cells expressing WT or mutant RyR1 with R-CEPIA1er. (**A**) Typical results of time-lapse R-CEPIA1er fluorescence measurement of WT (left), W882A (center), and R1000A (right) RyR1 cells. To activate RyR1, the assay media was supplemented with 5 mM caffeine. Dantrolene (0-10 μM) was added after 60 seconds (arrow). (**B**) Dose-dependent effect of dantrolene on [Ca^2+^]_ER_ in WT (black), W882A (red), W996A (green) and R1000A (blue) RyR1 cells. (**C)** IC_50_ values of dantrolene for WT or mutant RyR1s. Note that effect of dantrolene was reduced (W996A and R1000A) or lost (W882A) in the mutants. Data are mean ± SD (n = 6). *, p<0.05 vs control. (**D-G**). Effect of dantrolene on [^3^H]ryanodine binding to microsomes from HEK293 cells expressing WT or mutant RyR1. (**D**) Ca^2+^-dependent [^3^H]ryanodine binding of WT and **(E)** W882A RyR1 in the absence (control, open circles) and presence (filled circles) of 10 μM dantrolene. *, p<0.05 vs control. (**F)** Dose-dependent effect of dantrolene at 30 μM Ca^2+^. (**G)** IC_50_ values for dantrolene. Note that dantrolene inhibition was reduced (W996A and R1000A) or lost (W882) in the mutants. Data are mean ± SD (n = 4). *, p<0.05 vs control.

Further, we tested the effect of dantrolene on the channel properties of RyR1 using [^3^H]ryanodine binding (*35*). Since ryanodine binds only to open RyR1s, ryanodine binding is a useful measure for the channel activity. Addition of 10 μM dantrolene to WT RyR1 significantly diminished the [^3^H]ryanodine binding at all Ca^2+^ concentrations (Fig. 2D). W882A RyR1 was insensitive to dantrolene (Fig. 2E). A concentration dependence curve of dantrolene at the maximally activating Ca^2+^ concentration revealed intermediate results for the W996A and R1000A mutations (Fig. 2F), where the IC_50_ for dantrolene increased more than one order of magnitude, and no measurable inhibition was detected on W882A RyR1 by dantrolene (Fig. 2G). Our functional studies support that the primary site identified using cryo-EM is responsible for its biological activity. An orientation of dantrolene such that π-π stacking interaction with W882 is crucial since the W882A mutant showed a complete abolishment of dantrolene inhibition (Fig. 2A, E). The other hydrophobic interaction with W996 is also important as seen by the >10-fold decrease in IC_50_ of dantrolene inhibition (Fig. 2B, F). Loss of charged interaction with R1000 resulted in ∼10-fold decrease in dantrolene inhibiting potency (Fig. 2C, G). Thus, all three of our validation mutants exhibited marked and substantial decrease in dantrolene inhibition supporting our proposed binding pose.

### Dantrolene reverses large-scale conformational changes induced by the Y523S mutation

The mutated residue, Y523, resides at the approximate center of an α-helical bundle in the N-terminal subdomain C (NTDC) (Fig. S1, inset). As elucidated by us previously (*29*), the Tyr-to-Ser mutation abolished the “spacer” role of Y523 inducing local rearrangement of the α helical bundle and triggering large-scale conformational changes that are transmitted across multiple domains. This consequence of the Y523S mutation causes the cytoplasmic assembly of the channels to adopt a pre-activated conformation (i.e., quasi-open) under closed-state conditions, thereby sensitizing the channels to more facile opening. When the displacement of domains of mutant RyR1 was measured and represented as a heat map, there was an obvious “cooling” (Figs. 3 A and S9), indicating that addition of dantrolene begins to reverse some of the large-scale conformational changes, stabilizing the channel in a “less primed”-like conformation as seen in the RyR1^YS^-closed-DAN map. A similar cooling effect was also observed for the RyR1^YS^-open-DAN dataset (Fig. 3B and S9). In both datasets, the biggest conformational change observed was at the P1 domain (Figs. 3 A-B and S9). The second-largest conformational change was undergone by the three sp1a Ryanodine (SPRY) domains (SPRY1, SPRY2 and SPRY3; residues 632-846 and 1056-1650), which neighbor the P1 domain. The magnitude of the observed changes decreased in magnitude radiating further away from the P1 domain (Figs. 3 A-B and 4). In the RyR1^YS^-open-DAN dataset, the reversal of conformational changes was more pronounced in domains further away from the P1 domain as compared to RyR1^YS^-closed-DAN and included CytA domains that were closer to the TmD such as the central domain (CD; residues 3668-4070) and its downstream domains, the EF hands and U motif (residues 4071-4131 and 4132-4251, respectively) (Figs. 3 A-B and S9), which appears to result from the additional effect of Ca^2+^ in the CD. Addition of dantrolene to the mutant channel did not entirely rescue the typical conformation of RyR1 in the closed and open states, respectively. In addition, the degree of overlap with the WT varied for different regions of the structure (Fig. S10), suggesting a distinct state.

**Figure 3.**
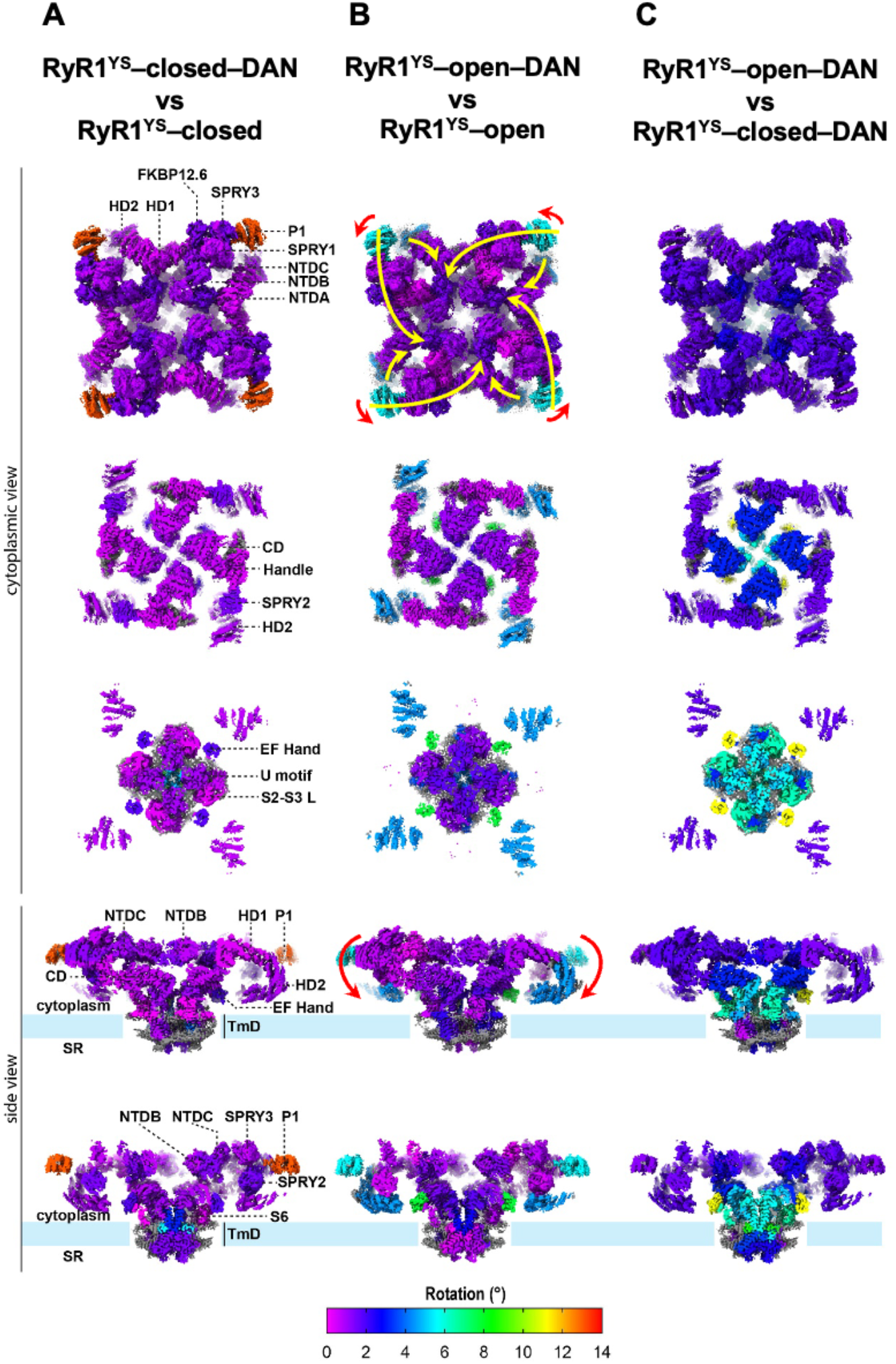
Heat map of movements across the various domains of RyR1 induced by addition of dantrolene to the Y523S MH/CCD mutant. **(A-B)** Rotations (in degrees) were measured in pair-wise comparison between the mutant and the mutant incubated with 50 μM dantrolene. For RyR1^YS^-EGTA and RyR1^YS^-Ca^2+^ we utilized our previously published models (PDB: 7T65 and PDB:7T64, respectively). The Y523S (YS) MH mutation induced a primed or quasi-open state of RyR1 under closed conditions, and an open^+^ conformation of RyR1, with larger magnitude of open-state movements (*29*). RyR1 domain names are indicated in the left panels. Arrows in (B) indicate main conformational changes induced by dantrolene in the presence of either EGTA or Ca^2+^. **(C)** Gating of mutant RyR1 in the presence of dantrolene displayed less movement of the peripheral domains (NTD, SPRY, handle, HD1, P1, HD2) and more similar movement of the central module (CD, EF hand, U motif, S6, CTD), when compared to gating of WT RyR1. The more tempered energetic landscape induced by dantrolene probably explains the lower probability of opening. See Fig. S9 for numeric information.

Binding of dantrolene caused a folding-in of the external arm of the V-shaped domain P1 by 10 Å, altering its center of mass by 3 Å and 13° (Fig. S9), inducing a movement of the SPRY complex and handle domain of that subunit towards the fourfold axis. The conformational change also resulted in slight separation of P1 from the helical domain 1 (HD1) of the neighboring subunit, allowing the HD1 domain to move towards the channel’s axis (arrows in Fig. 3B). This subtle movement transmits to the CD and the rest of the domains closer to the pore. A drooping and rotation of HD2 also appeared to push the base of the cytoplasmic shell inward. Parallel changes can be observed in the open state, starting from a similar translation of the P1 domain (11 Å shift of the external arm, change in center of mass by ∼3 Å, and smaller rotation of 6°). A larger movement of 4.4 Å in the case of HD2 (versus 1 Å in the closed state) was observed. The similar observations from two independent comparisons made at two extreme conformations of RyR1 suggest genuine changes induced by dantrolene. Overall, dantrolene acts as a negative allosteric modulator that pushes most domains towards the fourfold axis, stabilizing the closed state.

Comparison of the mutant open and closed channels in the presence of dantrolene indicates that the channel is still able to gate (Fig. 3C) albeit with more tempered movements than these observed for WT RyR1 (*29*): the peripheral domains (NTD, SPRY, handle, HD1, P1, HD2) rotated an average of 2° compared to an average of 6° in regular gating, the CTD and U-motif rotated an average of 5° compared to 7° in regular gating, while the CD and EF hand domain appeared to rotate as much as in WT RyR1 gating (see Discussion).

### Dantrolene shrinks the pore of RyR1^YS^ without hindering Ca^2+^ permeation

The TmD of RyR1 consists of 4 sets of S1-S6 helices arranged in such a way that the conduction pathway is lined by S6 helices (1 from each subunit) which continue as the C-terminal domain (CTD). Dantrolene did not affect the pore profile under closed state conditions (compare Fig. 5 A and C). Presence of activating Ca^2+^ causes approximation of the CTD and CD, whereby the CTD lifts and the CD is pulled down, allowing coordination of Ca^2+^ at their interface in a site composed of negatively charged residues, E3893 and E3967, along with Q3970 and H3895 (CD) and backbone carbonyl of T5001 (CTD). The upward movement of the CTD upon coordination of Ca^2+^ is one of the driving impetuses for separation of S6 helices and channel opening. In the RyR1^YS^-open-DAN map, we observe the upward movement of CTD along with the clearly visible density of the Ca^2+^ ion coordinated to the aforementioned residues (Fig. S6B). At both the primary constriction point as well as the secondary gate, I4933 and Q4937, respectively, the radius of the pore was narrowed by ∼1 Å (decrease in distance between Cα atoms) in the presence of dantrolene (compare Fig. 5 B and D), still sufficient to allow conduction of Ca^2+^. Sub-classification of both datasets into 4 classes indicated a uniformity in pore diameter among the channel population (Fig. S11).

**Figure 4.**
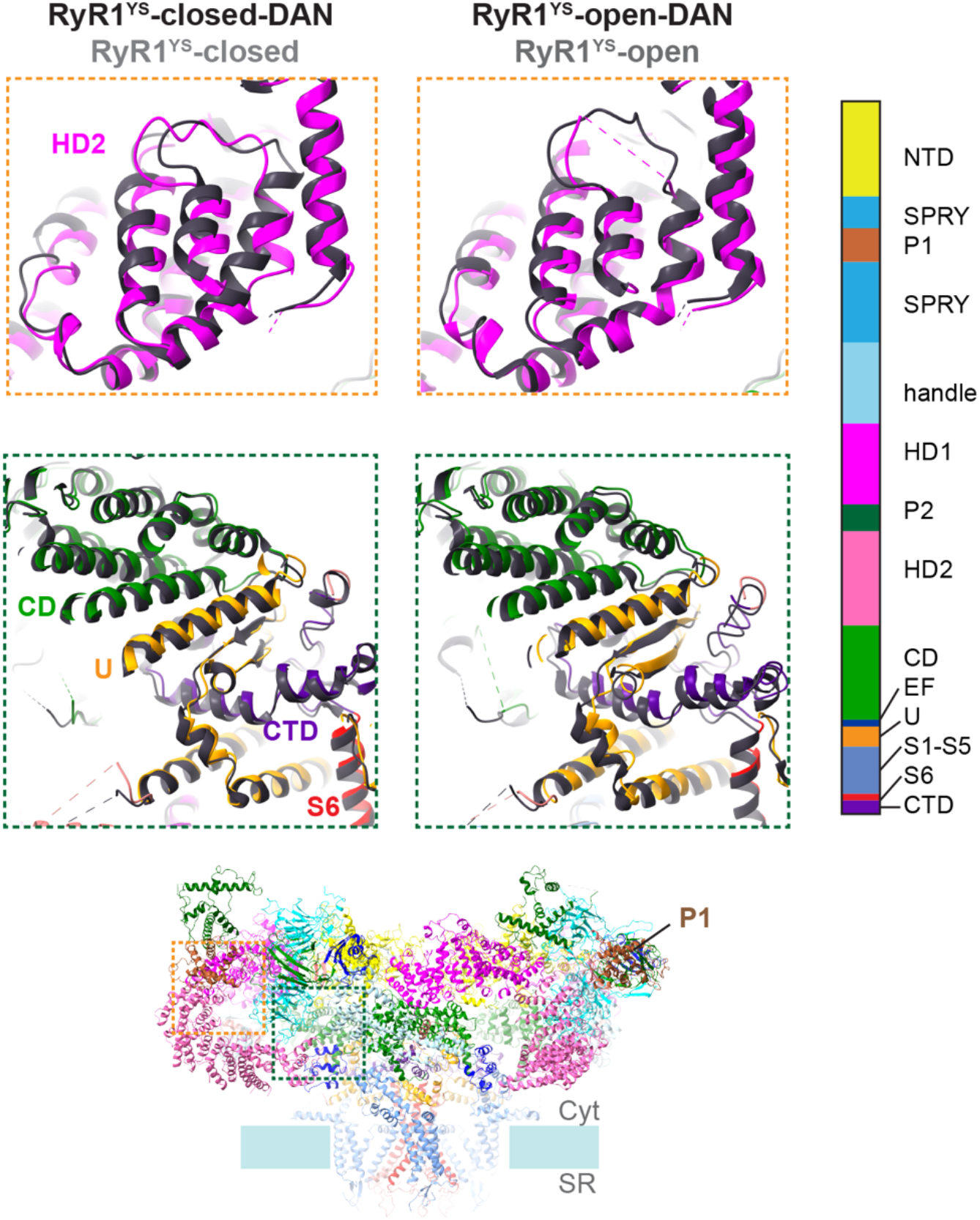
Binding of dantrolene triggers widespread conformational changes. The conformational changes, largest at the P1 domain, diminish in going from the peripheral (e.g. HD2), to the central (e.g. CD), to the transmembrane domain. Movements towards the fourfold axis are combined with rotational movements. The structure is color-coded according to domain and the mutant without dantrolene is shown in dark gray for reference.

**Figure 5.**
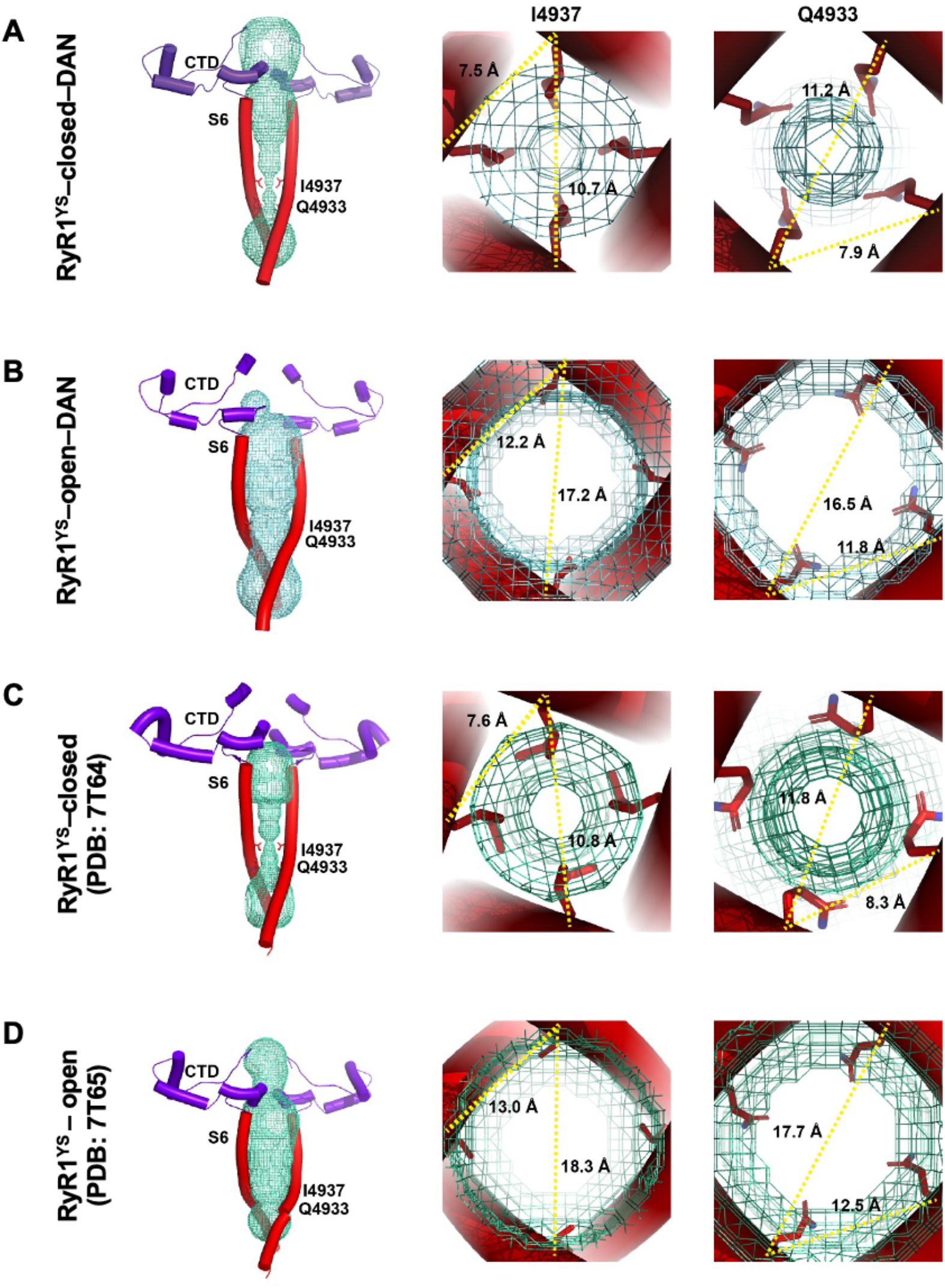
Pore Profiles of RyR1 Y523S in the presence of dantrolene under closed- and open-state conditions. Addition of dantrolene caused shrinking at both constriction points, I4937 and Q4933, by ∼1 Å under open-state conditions, which does not hinder Ca^2+^ permeation. The closed and open states for the mutant RyR1 are shown in the presence **(A-B)** and absence **(C-D)** of dantrolene.

## DISCUSSION

In recent years, there has been growing interest in discovery and development of drug-like molecules targeting disease mutant RyR1 channels (*35-38*). Despite this interest, dantrolene remains the only US FDA approved inhibitor of RyR1. Understanding how dantrolene inhibits RyR1 is crucial in designing better and newer analogs. Our cryo-EM structures show that dantrolene binds to the outermost four corners of the cytoplasmic domain of RyR1, nestled in the V shaped P1 domain. The π-π stacking interaction between the phenyl moiety of dantrolene and W882 plays a major role, with a secondary hydrophobic interaction with W996 and polar interactions with R1000. The proposed binding mode of dantrolene was validated with functional studies where interacting residues were mutated to alanine. Imaging the ER Ca^2+^ content in live cells showed that dantrolene reduced the constitutive Ca^2+^ leak of WT RyR1, and that the validation mutations curtailed the sensitivity to dantrolene, severely limiting its inhibitory effect. [^3^H]ryanodine binding data corresponded to the live-cell Ca^2+^ imaging results, where the validation mutants increased the IC_50_ for dantrolene. Mutating W882 entirely abolished any effect of dantrolene on RyR1, followed by reduction in potency by more than one order of magnitude for mutants W996A and R1000A in both assays. This indicates that dantrolene requires W882 for binding, and that W996 and R1000 are also critical. Among the two orientations possible, we favor the pose wherein the phenyl ring of dantrolene can form the crucial interaction with W882. Our determination of dantrolene’s binding site and its mode of binding should be very useful for future structure-guided design of analogs with increased inhibitory activity and greater water solubility.

Under open state conditions, dantrolene exerts a similar effect on P1 and similar conformational changes take place as in the closed state. In this case the orthosteric ligand Ca^2+^ overcomes the inhibitory effect of dantrolene and the channel opens, however the conformational change upon gating has different properties than WT RyR1. We compared the rotations and translations undergone by the mutant channel in the presence of dantrolene with previous measurements carried out on WT RyR1 using the same method (*29*). The individual rotations of the alpha helices in the transmembrane domain are practically the same as in WT RyR1, with special mention of the identical 8.5° rotation undergone by the S4-S5 linker, while overall translations are slightly smaller. In the CytA, most domains (NTD, SPRY, P1, handle, helical domains HD1 and HD2) undergo rotations and translations that are ∼32% and ∼34%, respectively, of these measured in WT RyR1 gating. The U-motif and CTD behave more similarly to WT (80% of the rotation and 55% of the translation undergone by WT RyR1 gating), while the CD and the EF hand domain undergo similar rotations and translations as WT RyR1. The CD/CTD and the EF hand domain contain the high and low affinity Ca^2+^ sensors, respectively (*34, 39*). These two sensors at the central module of the channel, in the interphase of the CytA and TmD, continue to control the pore, but their effect is dampened as dantrolene is bound to the peripheral module of the CytA. In summary, a clear picture emerges of Ca^2+^ as an orthosteric agonist while dantrolene acts as a negative allosteric modulator. The architecture of RyR1 permits gating but under a much more “controlling” peripheral CytA. By changing the energetic landscape, dantrolene affects the probability of opening but does not interfere with gating. Our results could explain in vivo studies demonstrating that gating proceeds, but the leak is much smaller in the presence of dantrolene (*10, 25*).

Dantrolene at clinical concentrations has been reported to have no effect on heart (which expresses the RyR2 isoform) (*24, 40*) or on recombinant RyR2 in HEK293 cells (*35*). This is in contrast with the high homology of the P1 domain between the RyR1 and RyR2 isoforms. Our structural results suggest two reasons that could account for the smaller response by RyR2. On one hand, the HD2 domain is consistently more flexible in the RyR2 reconstructions as discussed previously (*28, 41*); thus it is possible that the slight detachment of the helical domain from P1 noted in the presence of dantrolene has less repercussion for RyR2. On the other hand, the different quaternary arrangement of the two isoforms, with a most predominant corner-to-corner configuration in RyR1 than RyR2 (*42, 43*), could underlie an additional effect of dantrolene in RyR1 via an alteration of these quaternary interactions.

Overall, our structural and functional results provide important insight into dantrolene’s mode of action. The structural knowledge provided here will help develop dantrolene derivatives for anesthesia, spasticity, and potentially other RyR1-related disorders.

## MATERIALS AND METHODS

### Reagents

Reagents were purchased from Thermo Fisher Scientific or Sigma-Aldrich, with the exception of 1-palmitoyl-2-oleoyl-glycero-3-phosphocholine (POPC), membrane scaffold protein 1E3D1 (MSP1E3D1) and dantrolene, which were purchased from Addgene, Avanti Polar Lipids, and Tocris, respectively.

### Expression and purification of RyR1 Y523S from HEK293 cells

Recombinant rabbit RyR1 Y523S was expressed in and purified from human embryonic kidney (HEK) 293 cells as previously described (*29, 44*).

### Cell culture

Cell culture of HEK293 was performed on 150 mm plates using Dulbecco’s modified Eagle medium (high glucose, glutamine, and no sodium pyruvate) supplemented with fetal bovine serum and bovine growth serum (Gibco, Thermo Fisher Scientific) and penicillin-streptomycin. Expression of mutant channels was induced 48-72 h prior to harvesting by addition of doxycycline (∼1-2 μg/ml). Cells were collected by scrapping the plates were subjected to centrifugation (573 x g, 10 min). The resulting pellet was flash-frozen using liquid N_2_ and stored at -80 °C for further purification.

### Vesicle purification

Frozen cell pellet was thawed and ruptured by sonication (3 pulses of 20 sec each; 98% amplitude). The vesicle fraction of the ruptured cells was isolated by centrifugation (4,400 x g, 10 min) and subjected to ultracentrifugation (100,000 x g, 1 h) to pull down vesicles. The vesicle pellet was resuspended in 5 mM imidazole pH 7.4, 10% (w/v) sucrose, and protease inhibitor cocktail [aprotinin (5 μg/ml), leupeptin (5 μg/ml), and pefabloc (2.5 μg/ml)] (PI) using a Dounce homogenizer. Vesicles were flash-frozen using liquid _2_ and stored at -80 °C for subsequent protein purification.

### Protein purification

Frozen purified vesicles were solubilized for 25-30 min at 4 °C with mild agitation in solubilization buffer (5% CHAPS and 1.25% PC, 1 M NaCl, 20 mM Na-Mops pH 7.4, 2 mM DTT, and protease inhibitors (PI)). Insolubilized material were separated by ultracentrifugation (100,000 × g for 1 h). RyR1 Y523S was isolated using a sucrose density gradient. The supernatant from ultracentrifugation was layered on top of a discontinuous gradient composed of 6 layers – 10%, 12%, 14%, 16%, 18% and 20% w/v – and subjected to ultracentrifugation (120,000 x g, 23 h) using a swing bucket rotor. Fractions containing RyRs were pooled together and diluted 5-fold using dilution buffer (20 mM Na-Mops pH 7.4, 0.5% w/v CHAPS, 0.125% w/v PC, 2 mM DTT, and PI). Diluted RyRs were concentrated using a HiTrap Heparin HP column (1 mL; Amersham Biosciences) being subjected to two washings using wash buffer 1 (20 mM Na-Mops pH 7.4, 0.2 M NaCl, 0.5% w/v CHAPS, 0.125% w/v PC, 2 mM DTT, and PI) and wash buffer 2 (20 mM Na-Mops pH 7.4, 0.2 M NaCl, 0.015% w/v Tween-20, 2 mM DTT, and PI). Concentrated RyRs were eluted using elution buffer (20 mM Na Mops pH 7.4, 0.9 M NaCl, 0.015% w/v Tween-20, 2 mM DTT) in 100 μL aliquots and were stored at −80 °C after flash freezing using liquid N_2_. Sodium dodecyl sulfate-polyacrylamide gel electrophoresis (SDS-PAGE) and negative staining with 0.75% w/v uranyl formate were utilized to determine purity and quality of purified RyR1 Y523S. Frozen purified RyR1 Y523S were kept frozen until cryo-EM grid preparation.

### Reconstitution in nanodiscs

Purified rabbit RyR1 Y523S were reconstituted into nanodiscs as previously described (*29*). Briefly, purified RyR1 Y523S was mixed with MSP and POPC in a ratio of RyR1:MSP:POPC=1:2:50 and incubated at 4 °C for 3 h. This was followed by dialysis at 4 °C overnight to remove the detergent with the dialysis buffer being replaced after 2 h. Reconstituted RyR1 Y523S was incubated with adenosine 5’-triphosphate (ATP; 2 mM) and dantrolene (50 μM) for ∼1 h at 4 °C for both datasets. Additionally, ethylene glycol-bis(β-aminoethylether)-N,N,N’,N’-tetraacetic acid (EGTA) was added to both RyR1^YS^-closed-DAN (2 mM) and RyR1^YS^-open-DAN (0.5 mM) datasets prior to incubation. Stock solution of dantrolene (5 mM) was prepared by dissolving in DMSO at 50 °C over a period of 1 h followed by rotating at room temperature for 30 min. Free [Ca^2+^] for the open^+^–DAN was estimated to be ∼50 μM using Maxchelator (*45*). The quality and concentration of mutant channels was assessed by negative staining with 0.75% w/v uranyl formate and TEM imaging.

### Cryo-EM grid preparation

Cryo-EM grids were prepared as previously described (*29*). Sample (∼2.1 mg/ml) incubated with appropriate ligands was adsorbed onto Ultrafoil grids (R1.2/1.3, 300 mesh, Quantifoil) followed by blotting (blot force 1 to 2; blot time 1 to 2 s) and plunging into liquid ethane cooled by liquid N_2_ using a FEI Vitrobot Mark IV. No glow discharge was performed prior to adsorption. The chamber of the Vitrobot was maintained at a humidity of ∼95% and at 4 °C. Frozen grids were stored in liquid N_2_ for subsequent screening and data collection.

### Data collection

For both datasets, similar microscope parameters were utilized during data collection. Data was collected using a FEI Titan Krios equipped with a K3-Post GIF detector at an accelerating voltage of 300 kV, a magnification of 81,000 and pixel size of 1.08 Å using the software EPU (ThermoFisher Scientific). Dose rate corresponded to 50 e^-^/Å^2^ spread over 40 frames with a nominal defocus range of -1.2 to -2.6 μm. The slit width of the energy filter was set at 10 eV. See Table S1 for details.

### Image Processing

Steps in the image processing procedure for the two datasets, RyR1^YS^-DAN_closed and RyR1^YS^-open-DAN are shown in Figs. S1 and S2, respectively. For the RyR1^YS^-closed-DAN and RyR1^YS^-open-DAN datasets, 13,992 and 12,212 movies, respectively, were collected. Initial manual screening of movies resulted in removal of 452 and 625 movies for RyR1^YS^-closed-DAN and RyR1^YS^-open-DAN datasets, respectively, based on presence of crystalline ice, mistargeting, or streaking due to misalignment of energy filter. All image processing steps were performed using cryoSPARC (*46*) unless otherwise stated. The remainder of movies, 13,540 and 11,587 were subjected to motion correction while retaining all 40 frames and at the original pixel size of 1.08 Å. Patch CTF estimation was performed on motion corrected micrographs followed by curation to remove micrographs whose CTF fit resolution estimates and total full-frame motion were above 15 Å and 20 pixels, respectively. This resulted in elimination of 1,227 and 816 micrographs from the RyR1^YS^-closed-DAN and RyR1^YS^-open-DAN datasets, respectively. 2D reference classes generated from ∼2,000 particles were used to perform automated template-based particle picking on the final 12,313 and 11,396 micrographs of the RyR1^YS^-closed-DAN and RyR1^YS^-open-DAN datasets, respectively. The ∼1.7 and ∼1.9 million extracted automated picks were binned 4X to 4.32 Å to speed up subsequent rounds of 2D classification to remove false picks. Successive 4 and 5 rounds of 2D classifications (Figs. S1 and S2) resulted in 254,054 and 377,079 true particles retained for the RyR1^YS^-closed-DAN and RyR1^YS^-open-DAN datasets, respectively, and were further subjected to per-particle motion correction. For the RyR1^YS^-closed-DAN dataset 3D classification in to 3 classes segregated 249,034 particles into 1 good class, which upon further non-uniform refinement (*47*) with per-particle defocus and global CTF refinement applied yielded a final 3D reconstruction with a resolution of 3.34 Å (FSC_0.143_). For the RyR1^YS^-open-DAN dataset, the 377,079 true particles from 2D classification were further classified into 4 classes using ab-initio reconstruction. Approximately 50% of the 377,079 particles separated into 1 “good” class which were further subjected to 3D classification to give 189,847 final particles. The 3D reconstruction with a resolution of 3.23 Å (FSC_0.143_) was achieved using non-uniform refinement with per-particle defocus and global CTF refinement applied. To improve electron density around the less resolved P1 region encompassing the dantrolene binding site, local refinement was performed on the corner region consisting of SPRYs (1 to 3), P1, FKBP12.6 of one subunit and HD1, P2 and HD2 of adjoining subunit, reaching 3.8 and 3.2 Å resolution for the closed and open conformations, respectively. Composite maps were built with the focused corner domains. Masks utilized for particle subtraction and subsequent local refinement were generated using the molmap command on deposited structure of RyR1 Y523S (PDB ID: 7T64) in Chimera (*48*). Local resolution was determined using for the consensus maps as well as focused maps using cryoSPARC (*46*).

### Model building

A single subunit of previously published model of rabbit RyR1 Y523S under closed-state condition (PDB: 7T64) was used as the starting model except for region 1420-1625 which was modeled after PDB:8VJJ. Initial manual docking along with appropriate ligands (ATP, dantrolene, Ca^2+^) was performed on the composite map using Chimera (*48*). Model building was achieved by iterative rounds of real-space refinement with Ramachandran and secondary structure restraints applied in Phenix (*49*) followed by manual fitting using Coot (*50*). This iterative process of refinement and manual fitting was performed until optimum model quality was achieved as indicated by MolProbity scores (*51*).

### Analysis of structures

Individual domain movements (translation and rotation) were measured using PyMol (The PyMOL Molecular Graphics System, Version 2.5.2 Schrödinger, LLC) using the angle_between_domains script. Movement of the individual helices of the TmD was measured in Chimera (*48*) using the Axis/Planes/Centroids functionality. Rotation and translation were measured only for Cα atoms. Heat maps were generated by plugging in values of movement (rotation in °) in GraphPad Prism 9 (GraphPad Software, CA, USA) and projected on the cryo-EM maps using ChimeraX (*52*). Radius along the ion conduction pathway was measured using HOLE (*53*) and only the pore lining S6 helices and the CTD of the 4 subunits were retained. High resolution images for figures were generated using either Chimera (*48*), ChimeraX (*52*) or PyMol (The PyMOL Molecular Graphics System, Version 2.5.2 Schrödinger, LLC).

### Generation of stable RyR1 mutant cells

Stable HEK293 cells expressing mutant RyR1 induced by doxycycline were generated as described previously (*54*). Each validation mutation (W882A, W996A and R1000A) was introduced to rabbit RyR1 cDNA (GenBank accession number X15209.1) by inverse polymerase chain reaction using a BsiWI–BstBI fragment (pBS-RyR1cs3) from the cDNA cassettes encoding the full-length rabbit skeletal muscle RyR1 (pBS-RyR1) (*55*). The mutated fragment was then subcloned into an expression vector (pcDNA5/FRT/TO-RyR1). The expression vector was transfected into HEK293 cells, and clones with suitable RyR1 expression were selected and used for the experiments.

### Time-lapse ER Ca^2+^ measurements

Time-lapse ER Ca^2+^ measurements were performed as described previously (*35*). Briefly, HEK293 cells expressing WT or mutant RyR1 were seeded on 96-well, flat, clear-bottom black microplates (#3603; Corning, New York, NY) at a density of 2×10^4^ cells/well. One day after seeding, expression of RyR1 was induced by addition of doxycycline (2 μg/ml) to the culture medium. After 24 hours of induction, the culture medium was replaced with 81 μL of HEPES-buffered Krebs solution (140 mM NaCl, 5 mM KCl, 2 mM CaCl_2_, 1 mM MgCl_2_, 11 mM glucose, and 5 mM HEPES, pH 7.4), and the microplates were placed in the FlexStation3 fluorometer, which was preincubated at 37°C. R-CEPIA1er signals, which were excited at 560 nm and emitted at 610 nm, were captured every 10 seconds for 300 seconds. 51 μL of the dantrolene solution was applied to the cells 100 seconds after starting. The fluorescence change induced by the compounds was expressed as *F*/*F*_0_, in which averaged fluorescence intensity of the last 100 seconds (*F*) was normalized to that of the initial 100 seconds (*F*_0_).

### [^3^H]Ryanodine binding

[^3^H]Ryanodine binding was carried out as described (*35*). Microsomes isolated from the HEK293 cells were incubated for 2 h at 37°C with 5 nM [^3^H]ryanodine in a medium containing 0.17 M NaCl, 20 mM 3-(N-morpholino)-2-hydroxypropanesulfonic acid (MOPSO) at pH 7.0, 2 mM dithiothreitol, 1 mM AMPPCP and various concentrations of free Ca^2+^ buffered with 10 mM ethylene glycol-bis(2-aminoethylether)-N,N,N’,N’-tetraacetic acid (EGTA). Free Ca^2+^ concentrations were calculated using the WEBMAXC STANDARD (https://somapp.ucdmc.ucdavis.edu/pharmacology/bers/maxchelator/webmaxc/webmaxcS.htm) (*45*). [^3^H]Ryanodine-binding data (*B*) were normalized to the maximum number of functional channels (*B*_max_), which were separately determined by Scatchard plot analysis using various concentrations (3–20 nM) of [^3^H]ryanodine in a high-salt medium. The resultant *B*/*B*_max_ represents the average activity of each mutant.

## ACKNOWLEDGEMENTS

We are grateful to the members of the Radioisotope Research and Research Support Center of the Juntendo University Graduate School of Medicine for technical assistance. Cryo-EM grid preparation, initial screening and data collection was carried out at University of Virginia cryo-EM facility (supported by S10-RR025067). Funded by National Institute of Health grant R01 AR068431, U24 GM116789, NIH U24 GM116790 (to MS), American Heart Association Post-Doctoral Fellowship 19POST34430178 (to KAI), Japan Society for the Promotion of Science grant 19H03404 and 22H02805, Japan National Center of Neurology and Psychiatry (5-6), Research Center for Biomedical Engineering (4001) and The Naito Foundation (to TM).

## AUTHOR CONTRIBUTIONS

Conceptualization: MS, Formal Analysis: KAI, Funding Acquisition: KAI, TM, MS; Investigation: KAI, TK, TM, MS, Supervision: MS, Visualization: KAI, MS, Writing – original draft: KAI, MS; Writing – review & editing: KAI, TM, MS.

## DATA AND CODE AVAILABILITY

All data are available in the main text or the supplementary materials. Cryo-EM composite maps (full map combined with P1-focused map) have been deposited in Electron Microscopy Data Bank (EMDB), www.emdatabank.org, with the following IDs: EMD-45584 for the RyR1^YS^–DAN-closed and EMD-45585 for the RyR1^YS^-open-DAN datasets.

Cryo-EM unmodified maps have been deposited in Electron Microscopy Data Bank (EMDB) with the following IDs: EMD-45493 for the RyR1^YS^–DAN-closed and EMD-45497 for the RyR1^YS^-open-DAN datasets.

Atomic models for the two datasets based on the composite map have been deposited in the Protein Data Bank (PDB), www.pdb.org, with the following IDs: EMD-9CGP for the RyR1^YS^– DAN-closed and EMD-9CGQ for the RyR1^YS^-open-DAN datasets.

## REFERENCES

1. K. A. Iyer, V. Barnakov, M. Samso, Three-dimensional perspective on ryanodine receptor mutations causing skeletal and cardiac muscle-related diseases. Curr Opin Pharmacol. 68, 102327 (2023).

2. T. A. Lawal et al., Ryanodine receptor 1-related disorders: an historical perspective and proposal for a unified nomenclature. Skelet Muscle 10, 32 (2020).

3. V. Bauerova-Hlinkova, D. Hajduchova, J. A. Bauer, Structure and Function of the Human Ryanodine Receptors and Their Association with Myopathies-Present State, Challenges, and Perspectives. Molecules 25, (2020).

4. H. Jungbluth, Central core disease. Orphanet J Rare Dis 2, 25 (2007).

5. L. Yang, T. Tautz, S. Zhang, A. Fomina, H. Liu, The current status of malignant hyperthermia. J Biomed Res 34, 75–85 (2019).

6. J. R. Lopez, L. Alamo, C. Caputo, J. Wikinski, D. Ledezma, Intracellular ionized calcium concentration in muscles from humans with malignant hyperthermia. Muscle Nerve 8, 355–358 (1985).

7. T. Yang et al., Pharmacologic and functional characterization of malignant hyperthermia in the R163C RyR1 knock-in mouse. Anesthesiology 105, 1164–1175 (2006).

8. J. Tong, T. V. McCarthy, D. H. MacLennan, Measurement of resting cytosolic Ca2+ concentrations and Ca2+ store size in HEK-293 cells transfected with malignant hyperthermia or central core disease mutant Ca2+ release channels. J Biol Chem 274, 693–702 (1999).

9. J. M. Eltit et al., RyR1-mediated Ca2+ leak and Ca2+ entry determine resting intracellular Ca2+ in skeletal myotubes. J Biol Chem 285, 13781–13787 (2010).

10. J. M. Eltit, X. Ding, I. N. Pessah, P. D. Allen, J. R. Lopez, Nonspecific sarcolemmal cation channels are critical for the pathogenesis of malignant hyperthermia. FASEB J 27, 991–1000 (2013).

11. J. R. Lopez, P. D. Allen, L. Alamo, D. Jones, F. A. Sreter, Myoplasmic free [Ca2+] during a malignant hyperthermia episode in swine. Muscle & Nerve 11, 82–88 (1988).

12. W. J. Durham et al., RyR1 S-nitrosylation underlies environmental heat stroke and sudden death in Y522S RyR1 knockin mice. Cell 133, 53–65 (2008).

13. J. Balderas-Villalobos, T. W. E. Steele, J. M. Eltit, Physiological and Pathological Relevance of Selective and Nonselective Ca(2+) Channels in Skeletal and Cardiac Muscle. Adv Exp Med Biol 1349, 225–247 (2021).

14. M. H. Dykes, Evaluation of a muscle relaxant: dantrolene sodium (Dantrium). JAMA 231, 862–864 (1975).

15. G. A. Gronert, J. H. Milde, R. A. Theye, Dantrolene in porcine malignant hyperthermia. Anesthesiology 44, 488–495 (1976).

16. G. G. Harrison, Control of the malignant hyperpyrexic syndrome in MHS swine by dantrolene sodium. Br J Anaesth 47, 62–65 (1975).

17. I. L. Anderson, E. W. Jones, Porcine malignant hyperthermia: effect of dantrolene sodium on in-vitro halothane-induced contraction of susceptible muscle. Anesthesiology 44, 57–61 (1976).

18. M. E. Kolb, M. L. Horne, R. Martz, Dantrolene in human malignant hyperthermia. Anesthesiology 56, 254–262 (1982).

19. T. Krause, M. U. Gerbershagen, M. Fiege, R. Weisshorn, F. Wappler, Dantrolene--a review of its pharmacology, therapeutic use and new developments. Anaesthesia 59, 364–373 (2004).

20. P. M. Hopkins et al., Malignant hyperthermia 2020: Guideline from the Association of Anaesthetists. Anaesthesia 76, 655–664 (2021).

21. FDA. (2014).

22. A. Ward, M. O. Chaffman, E. M. Sorkin, Dantrolene. A review of its pharmacodynamic and pharmacokinetic properties and therapeutic use in malignant hyperthermia, the neuroleptic malignant syndrome and an update of its use in muscle spasticity. Drugs 32, 130–168 (1986).

23. K. Paul-Pletzer, S. S. Palnitkar, L. S. Jimenez, H. Morimoto, J. Parness, The skeletal muscle ryanodine receptor identified as a molecular target of [3H]azidodantrolene by photoaffinity labeling. Biochemistry 40, 531–542 (2001).

24. F. Zhao, P. Li, S. R. Chen, C. F. Louis, B. R. Fruen, Dantrolene inhibition of ryanodine receptor Ca2+ release channels. Molecular mechanism and isoform selectivity. J Biol Chem 276, 13810–13816 (2001).

25. G. Cherednichenko et al., Enhanced excitation-coupled calcium entry in myotubes expressing malignant hyperthermia mutation R163C is attenuated by dantrolene. Mol Pharmacol 73, 1203–1212 (2008).

26. K. Paul-Pletzer et al., Identification of a dantrolene-binding sequence on the skeletal muscle ryanodine receptor. J Biol Chem 277, 34918–34923 (2002).

27. R. Wang et al., Localization of the dantrolene-binding sequence near the FK506-binding protein-binding site in the three-dimensional structure of the ryanodine receptor. J Biol Chem 286, 12202–12212 (2011).

28. K. A. Iyer et al., Structural mechanism of two gain-of-function cardiac and skeletal RyR mutations at an equivalent site by cryo-EM. Science advances 6, eabb2964 (2020).

29. K. A. Iyer, Y. Hu, T. Klose, T. Murayama, M. Samso, Molecular mechanism of the severe MH/CCD mutation Y522S in skeletal ryanodine receptor (RyR1) by cryo-EM. Proc Natl Acad Sci U S A 119, e2122140119 (2022).

30. K. A. Woll, O. Haji-Ghassemi, F. Van Petegem, Pathological conformations of disease mutant Ryanodine Receptors revealed by cryo-EM. Nat Commun 12, 807 (2021).

31. T. Murayama et al., Divergent Activity Profiles of Type 1 Ryanodine Receptor Channels Carrying Malignant Hyperthermia and Central Core Disease Mutations in the Amino-Terminal Region. PLoS One 10, e0130606 (2015).

32. M. Samso, A guide to the 3D structure of the ryanodine receptor type 1 by cryoEM. Protein Sci 26, 52–68 (2017).

33. S. Ducreux et al., Functional properties of ryanodine receptors carrying three amino acid substitutions identified in patients affected by multi-minicore disease and central core disease, expressed in immortalized lymphocytes. Biochem J 395, 259–266 (2006).

34. A. des Georges et al., Structural Basis for Gating and Activation of RyR1. Cell 167, 145–157 e117 (2016).

35. T. Murayama et al., Efficient High-Throughput Screening by Endoplasmic Reticulum Ca(2+) Measurement to Identify Inhibitors of Ryanodine Receptor Ca(2+)-Release Channels. Mol Pharmacol 94, 722–730 (2018).

36. T. Yamazawa et al., A novel RyR1-selective inhibitor prevents and rescues sudden death in mouse models of malignant hyperthermia and heat stroke. Nat Commun 12, 4293 (2021).

37. R. T. Rebbeck et al., RyR1-targeted drug discovery pipeline integrating FRET-based high-throughput screening and human myofiber dynamic Ca(2+) assays. Sci Rep 10, 1791 (2020).

38. A. Kushnir et al., Intracellular calcium leak as a therapeutic target for RYR1-related myopathies. Acta Neuropathol 139, 1089–1104 (2020).

39. A. R. Nayak, M. Samso, Ca(2+)-inactivation of the mammalian ryanodine receptor type 1 in a lipidic environment revealed by cryo-EM. Elife 11, e75568 (2022).

40. B. R. Fruen, J. R. Mickelson, C. F. Louis, Dantrolene inhibition of sarcoplasmic reticulum Ca2+ release by direct and specific action at skeletal muscle ryanodine receptors. J Biol Chem 272, 26965–26971 (1997).

41. S. Dhindwal et al., A cryo-EM-based model of phosphorylation- and FKBP12.6-mediated allosterism of the cardiac ryanodine receptor. Sci Signal 10, (2017).

42. C. Paolini, F. Protasi, C. Franzini-Armstrong, The relative position of RyR feet and DHPR tetrads in skeletal muscle. J Mol Biol 342, 145–153 (2004).

43. V. Cabra, T. Murayama, M. Samso, Ultrastructural Analysis of Self-Associated RyR2s. Biophys J 110, 2651–2662 (2016).

44. Y. Hu et al., Purification of Recombinant Wild Type and Mutant Ryanodine Receptors Expressed in HEK293 Cells. Bio-Protocol 11, (2021).

45. D. M. Bers, C. W. Patton, R. Nuccitelli, A practical guide to the preparation of Ca(2+) buffers. Methods Cell Biol 99, 1–26 (2010).

46. A. Punjani, J. L. Rubinstein, D. J. Fleet, M. A. Brubaker, cryoSPARC: algorithms for rapid unsupervised cryo-EM structure determination. Nat Methods 14, 290–296 (2017).

47. A. Punjani, H. Zhang, D. J. Fleet, Non-uniform refinement: adaptive regularization improves single-particle cryo-EM reconstruction. Nat Methods 17, 1214–1221 (2020).

48. E. F. Pettersen et al., UCSF Chimera—a visualization system for exploratory research and analysis. Journal of computational chemistry 25, 1605–1612 (2004).

49. D. Liebschner et al., Macromolecular structure determination using X-rays, neutrons and electrons: recent developments in Phenix. Acta Crystallogr D Struct Biol 75, 861–877 (2019).

50. P. Emsley, B. Lohkamp, W. G. Scott, K. Cowtan, Features and development of Coot. Acta Crystallogr D Biol Crystallogr 66, 486–501 (2010).

51. C. J. Williams et al., MolProbity: More and better reference data for improved all-atom structure validation. Protein Science 27, 293–315 (2018).

52. E. F. Pettersen et al., UCSF ChimeraX: Structure visualization for researchers, educators, and developers. Protein Sci 30, 70–82 (2021).

53. O. S. Smart, J. G. Neduvelil, X. Wang, B. A. Wallace, M. S. Sansom, HOLE: a program for the analysis of the pore dimensions of ion channel structural models. J Mol Graph 14, 354-360, 376 (1996).

54. T. Murayama et al., Genotype-Phenotype Correlations of Malignant Hyperthermia and Central Core Disease Mutations in the Central Region of the RYR1 Channel. Hum Mutat 37, 1231–1241 (2016).

55. J. Tong et al., Caffeine and halothane sensitivity of intracellular Ca2+ release is altered by 15 calcium release channel (ryanodine receptor) mutations associated with malignant hyperthermia and/or central core disease. J Biol Chem 272, 26332–26339 (1997).

