## Supplementary Material for "Dantrolene inhibition of ryanodine receptor 1 carrying the severe malignant hyperthermia mutation Y522S visualized by cryo-EM"

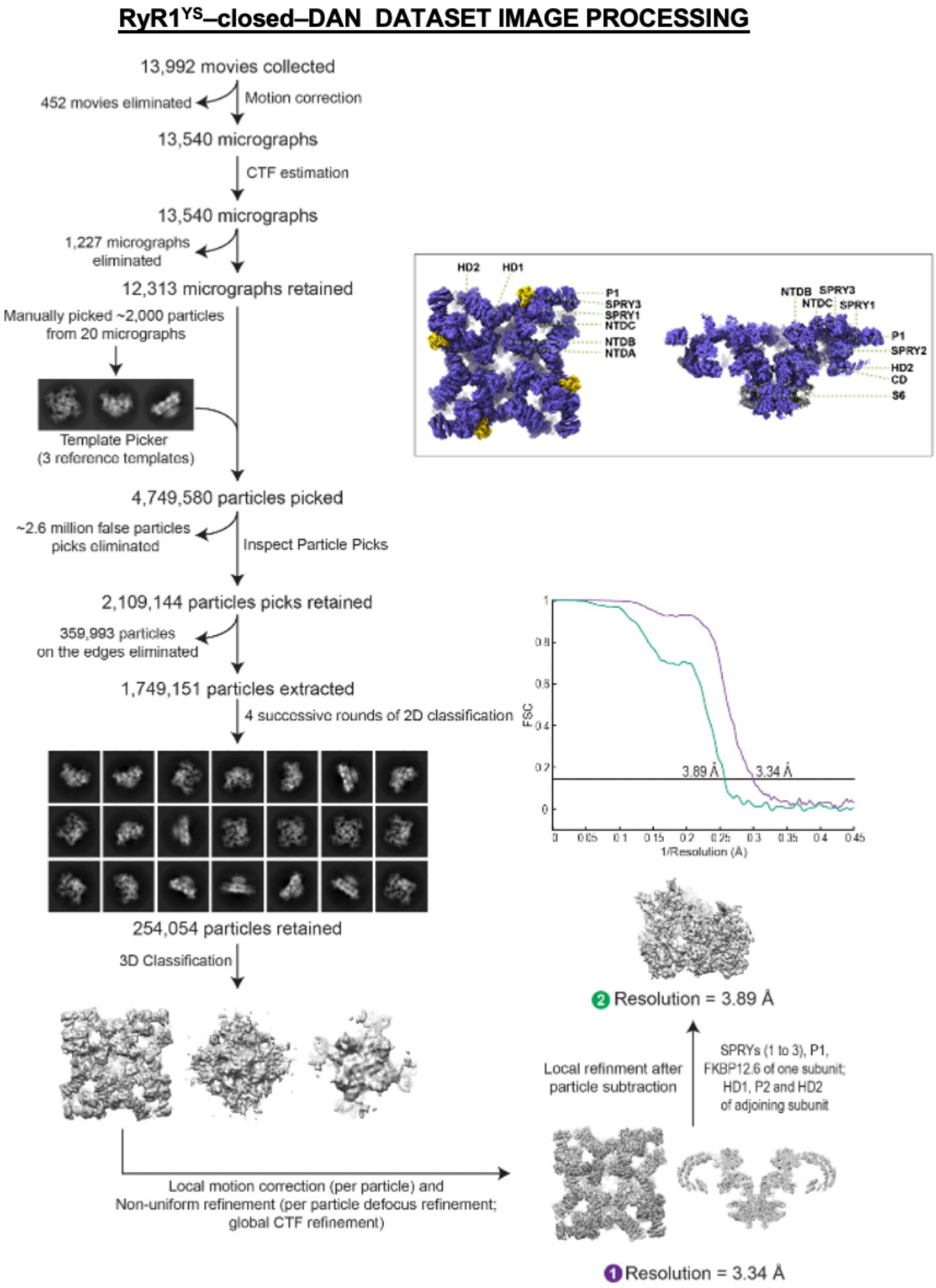

**Figure S1. Flowchart detailing steps involved in single-particle image processing of the RyR1$^{YS}$-closed-DAN dataset**

The mutant RyR1 with FKBP12.6 was prepared in the presence of ATP and EGTA and the transmembrane domain was embedded in nanodisc. The inset indicates the position of the different domains of the protein.

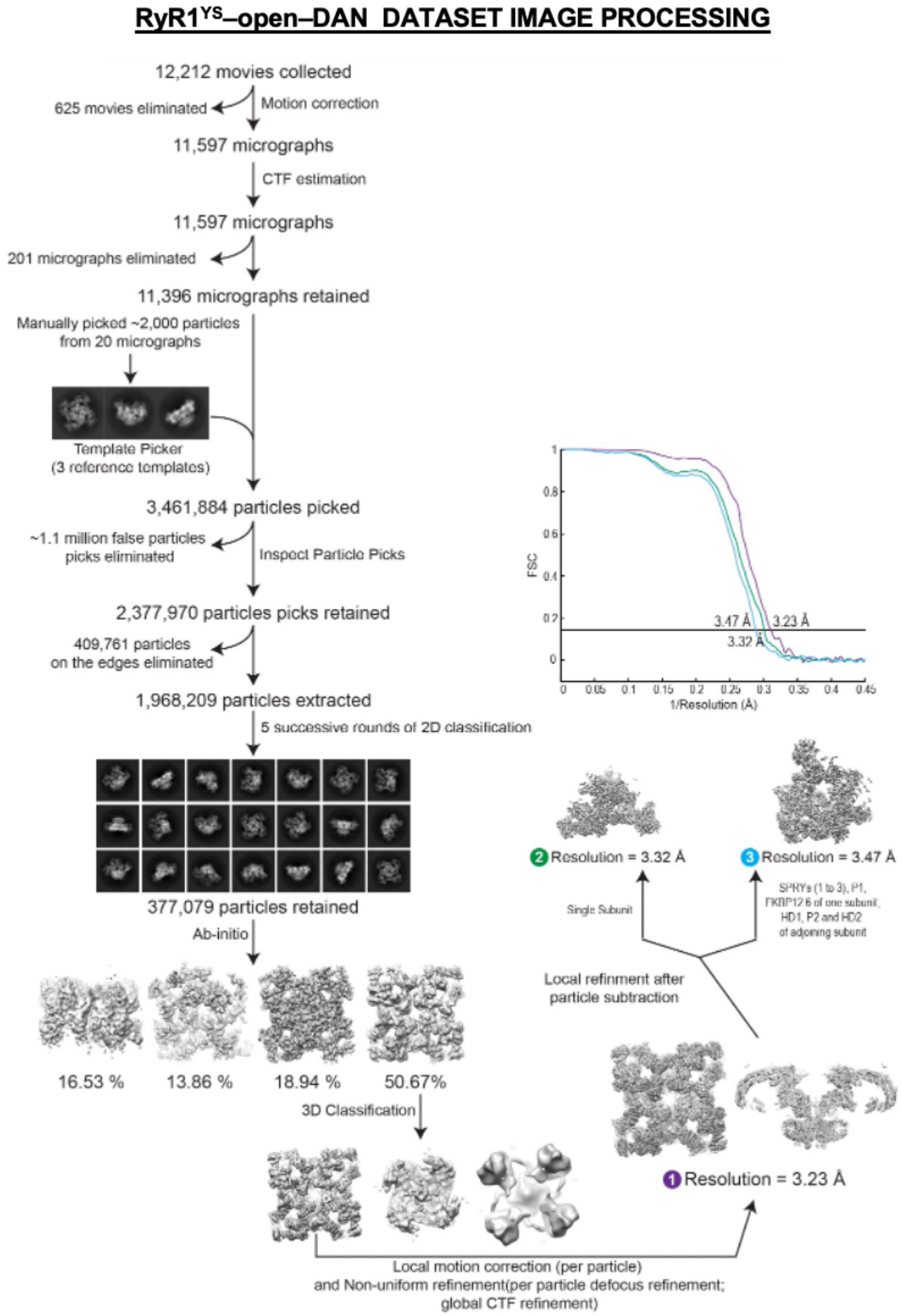

**Figure S2. Flowchart detailing steps involved in single-particle image processing of the RyR1$^{YS}$- open-DAN dataset**

The mutant RyR1 with FKBP12.6 was prepared in the presence of ATP and 50 μM free $Ca^{2+}$ and the transmembrane domain was embedded in nanodisc.

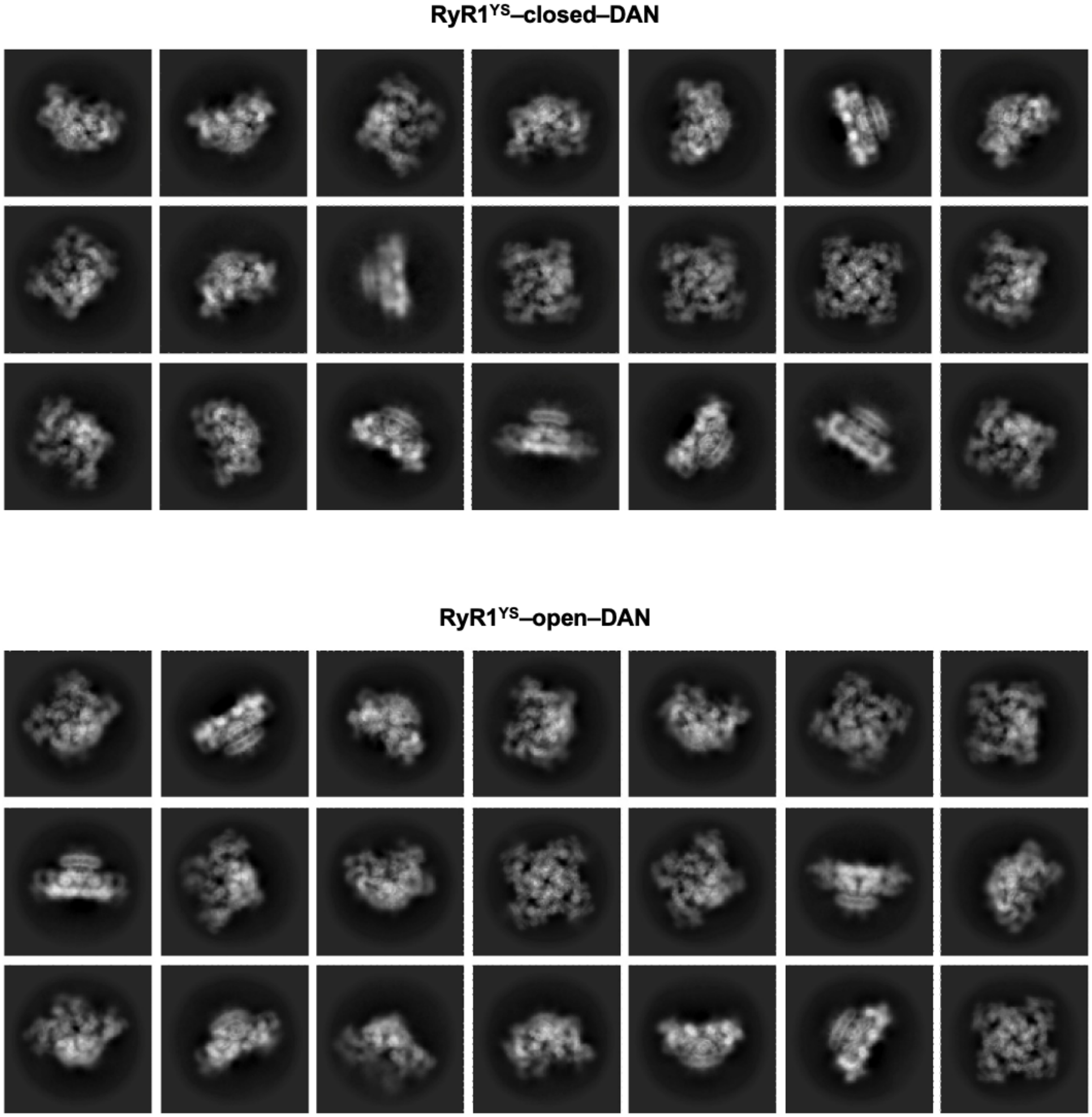

**Figure S3. 2D class averages for the datasets corresponding to RyR1$^{YS}$-closed (ATP/EGTA) with dantrolene and RyR1$^{YS}$-open (ATP/$Ca^{2+}$) with dantrolene**

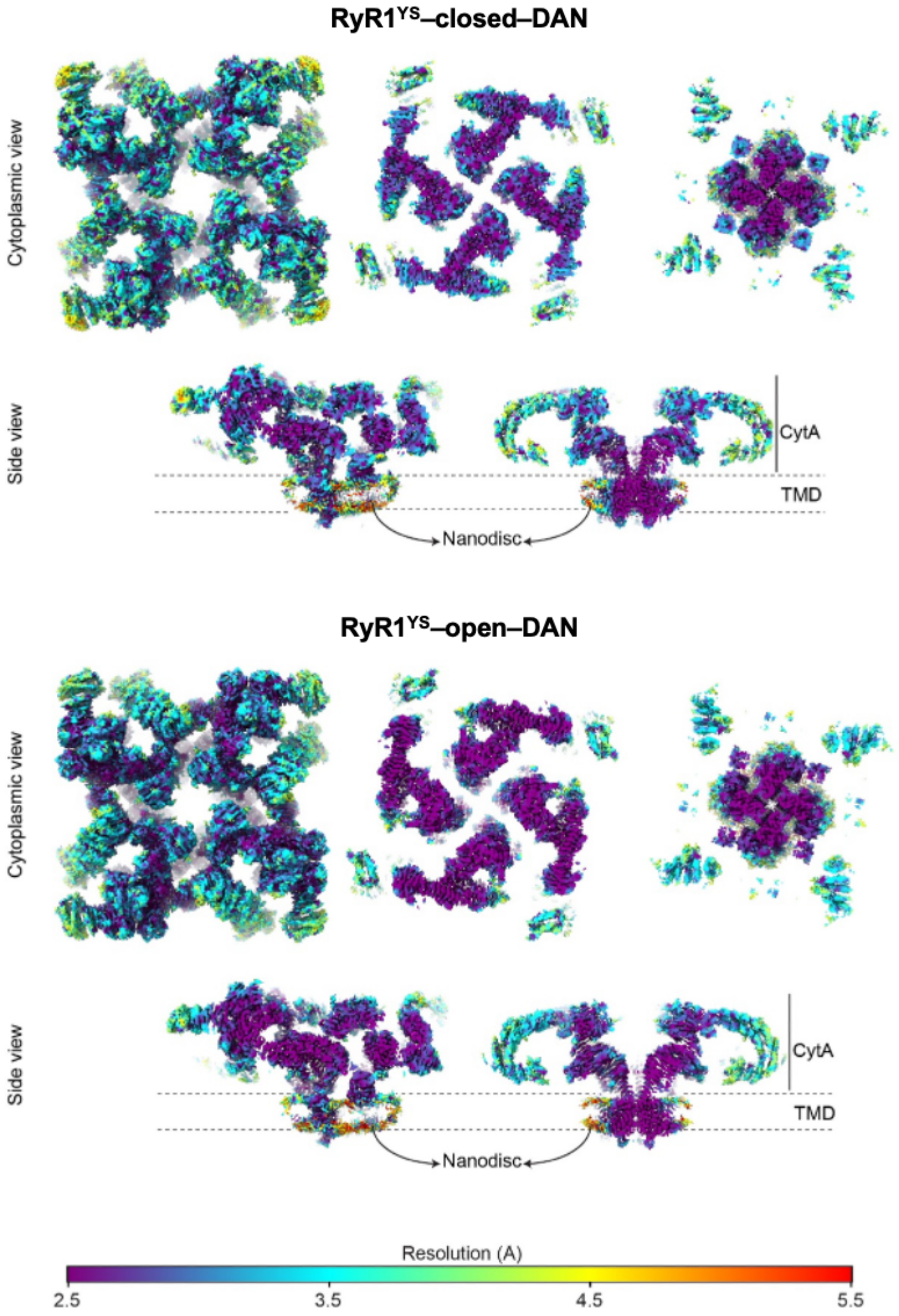

**Figure S4. Cryo-EM maps of the RyR1$^{YS}$-EGTA with dantrolene and RyR1$^{YS}$-$Ca^{2+}$ with dantrolene color-coded according to local resolution**

Maps correspond to the original structures. The resolution of the corner domains was further improved (to 3.8 Å and 3.2 Å for the closed and open reconstructions, respectively) using a focused mask.

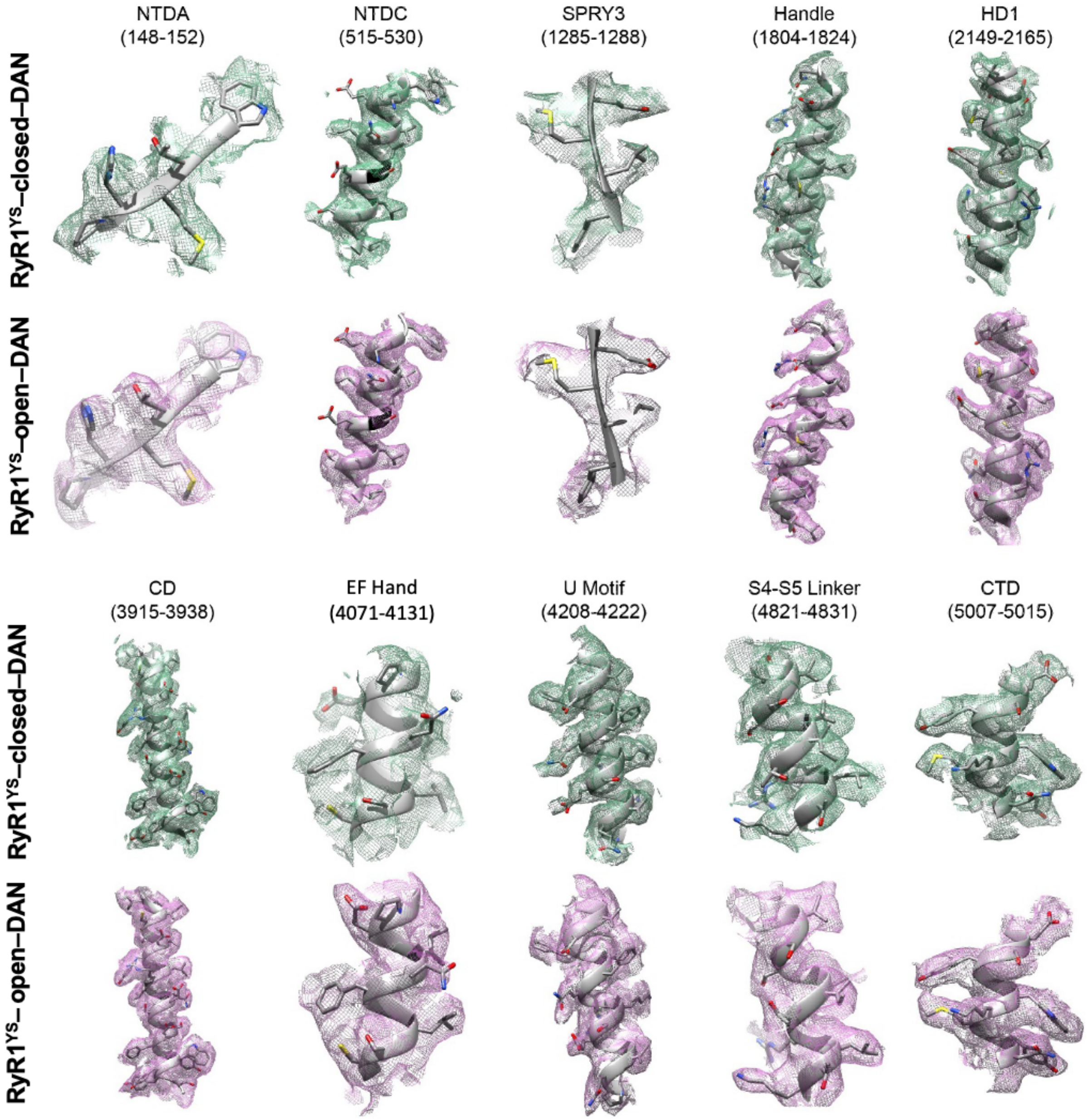

**Figure S5. Visualization of side chains in different regions of the 3D reconstruction**

Both the RyR1 Y523S cryo-EM maps (green and pink mesh, respectively) obtained in the presence of dantrolene inhibitor under closed and open state conditions show secondary structure details with clarity, allowing modeling of side chains of residues with ease.

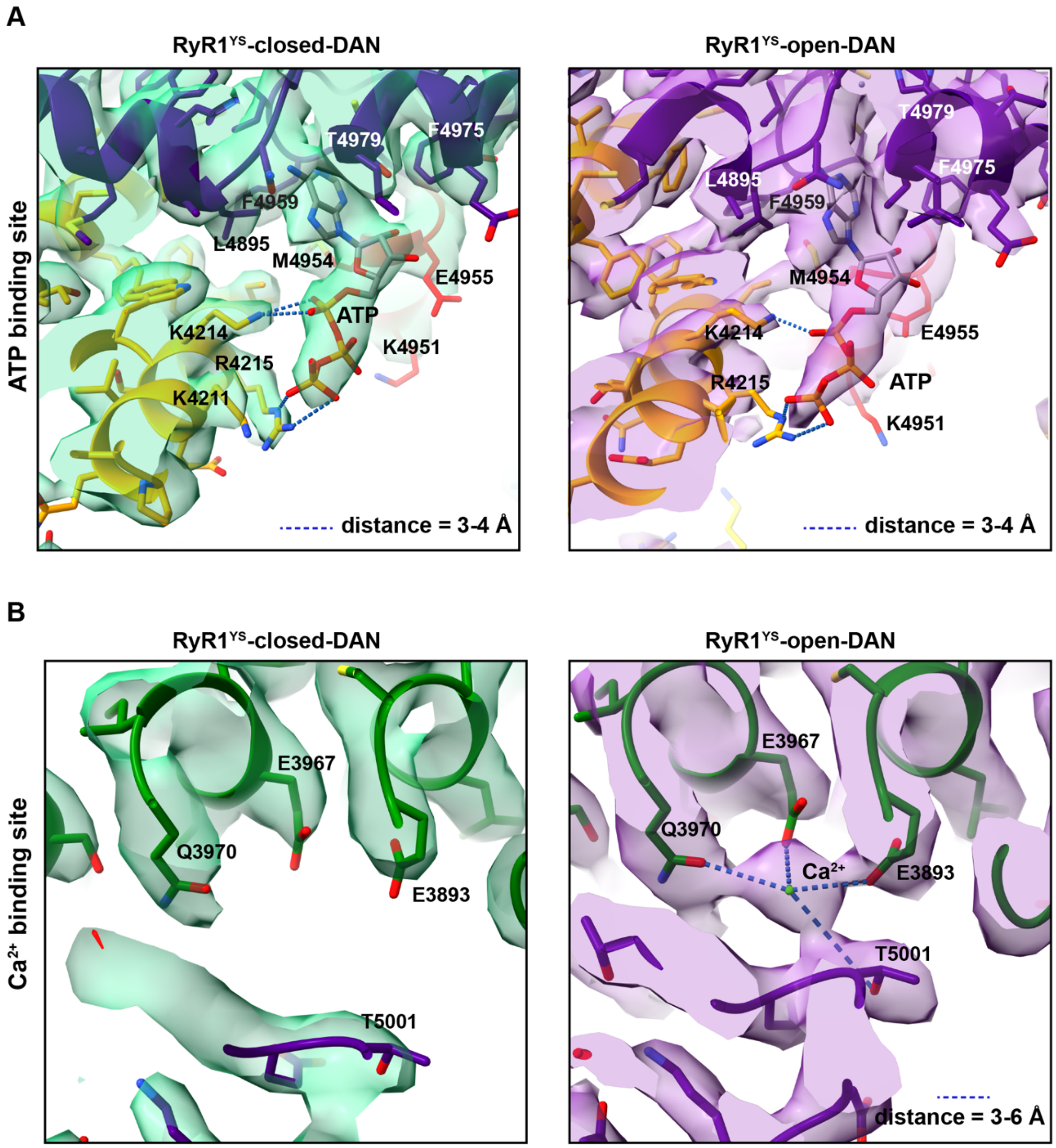

**Figure S6. Ligand binding sites in RyR1**

**(A)** Cryo-EM map of RyR1 with occupied ATP binding site. ATP interacted with residues at the interface of S6 (red), U-motif (yellow) and CTD (indigo) of the same subunit under both closed- and open-state conditions. **(B)** Cryo-EM map of RyR1 showing the $Ca^{2+}$ binding site, formed by the CD (green) and CTD (indigo) domains. Density for $Ca^{2+}$ was visible only in the $RyR1^{YS}$-open-DAN cryo-EM map.

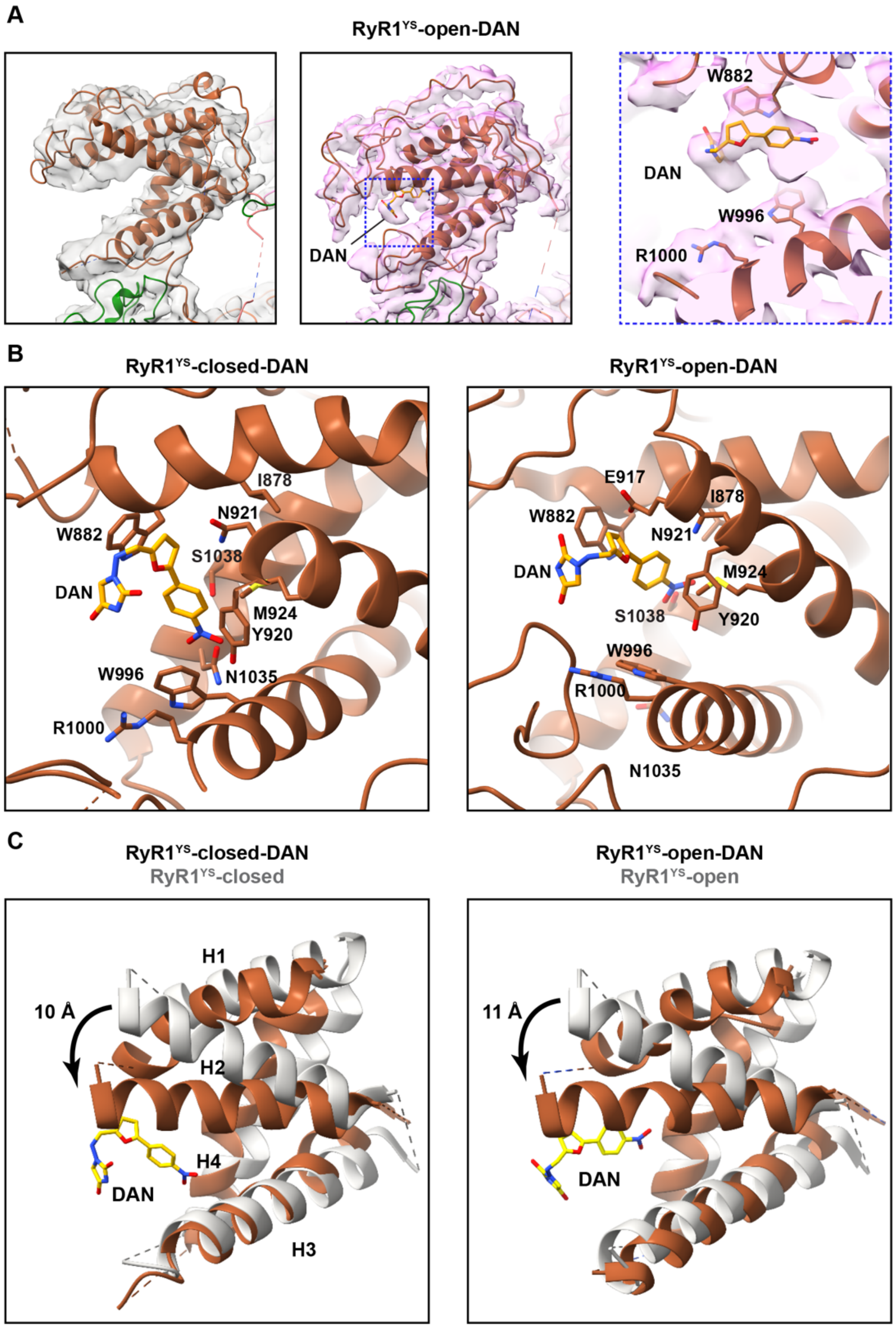

**Figure S7**. **Detail of the dantrolene binding site and its effect on the P1 domain**

(**A**) Conformational changes induced by dantrolene on the P1 domain for RyR1$^{YS}$-Ca$^{2+}$ (open state)**.** The cryo-EM density with validated residues is shown on the right also for the open state. (**B**) Residues in the P1 crevice within interacting distance of dantrolene are highlighted. Orientation of each panel is optimized to display residues of interest. (**C**) The position of the P1 alpha helices changes upon addition of dantrolene, where H1 and H2 are pulled toward H3 and H4. The position of the helices without dantrolene is shown in gray for comparison.

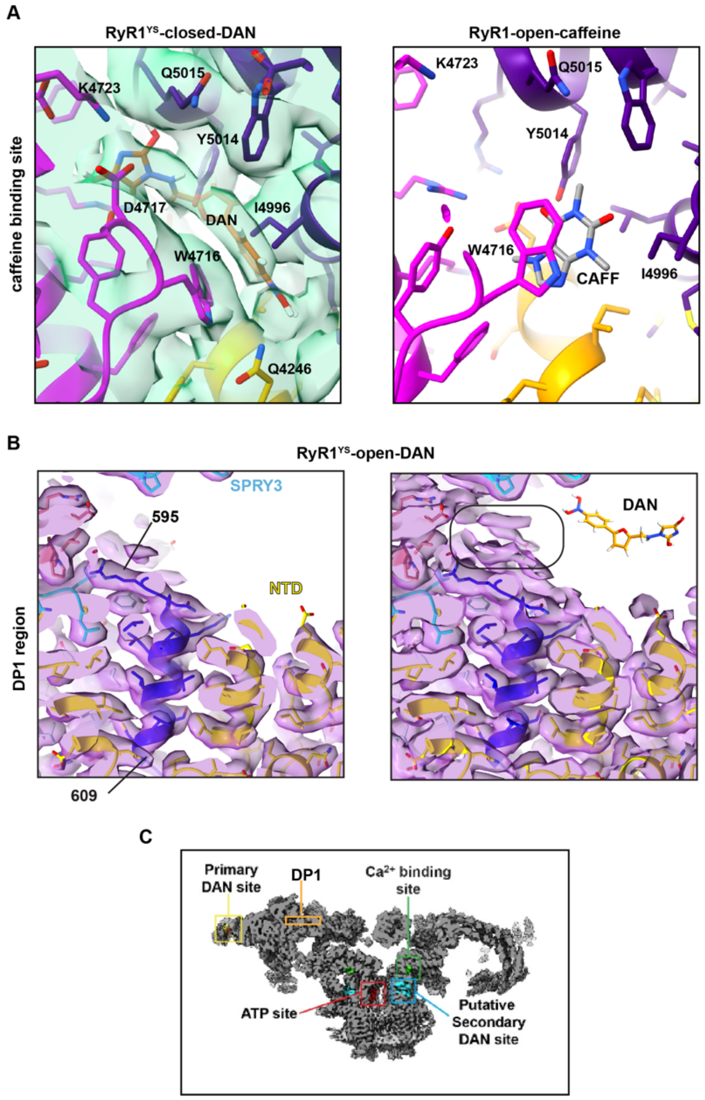

**Figure S8. Putative secondary binding site for dantrolene and postulated DP1 binding site**

**(A)** Partial density was observed at the interface of CTD (indigo) and U-motif (yellow) of one subunit and S2-S3 linker (magenta) of the neighboring subunit. This putative secondary dantrolene binding site was observed under the closed-state condition only and is near the previously identified caffeine binding site observed in the presence of $Ca^{2+}$ and caffeine (WT RyR1 open-state; PDB: 5TAL) shown on the right for comparison. (**B**) Region of the NTD (yellow) containing the DP1 peptide (blue), which binds dantrolene as a synthetic peptide *(26).* The cryo-EM density around the DP1 peptide was fully accounted by side chains (left panel). Although extra cryo-EM densities in the empty space near residues 595-596 appear at a lower threshold (right panel, rounded rectangle), they are too small to fit the dantrolene molecule (shown to scale). (**C**) Side view of RyR1 showing the positions of the primary dantrolene binding site, ligand binding sites, putative secondary binding site, and DP1 sequence.

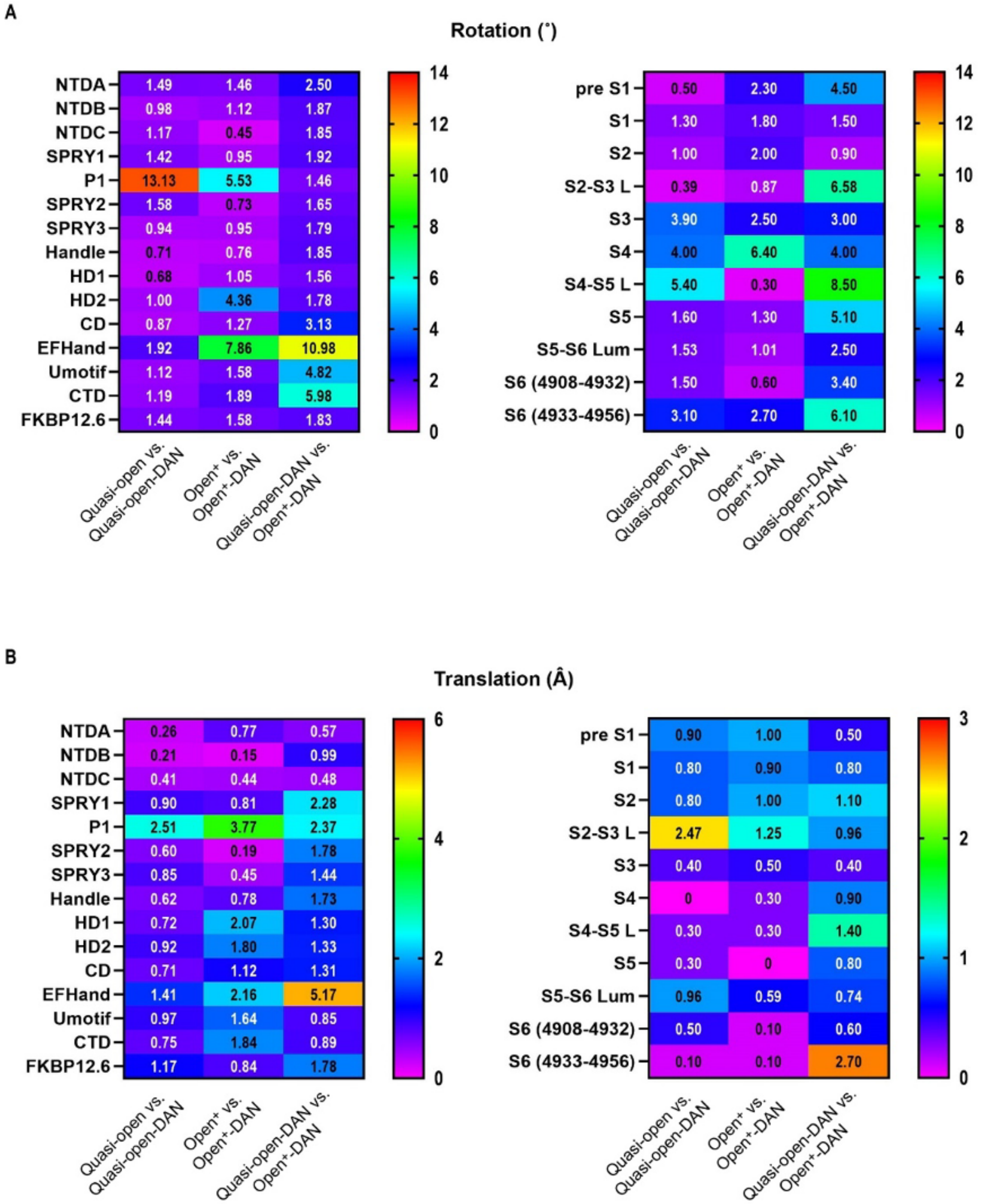

**Figure S9. Quantification of the conformational reversion caused by dantrolene**

(**A** and **B**) Values for individual domain movements (rotation in ° and translation in Å) corresponding to the heat map. Calculations were performed in pair-wise comparison in Pymol with respect to our previously published models for $RyR1^{YS}$ prepared under closed (PDB: 7T64) and open-state conditions (PDB IDs: 7T64 and 7T65, respectively). The terms quasi-open and $open^{+}$ describe the anomalous conformations of the channel with the gain-of-function mutation. The columns on the right measure the gating movement of the mutant channel in the presence of dantrolene.

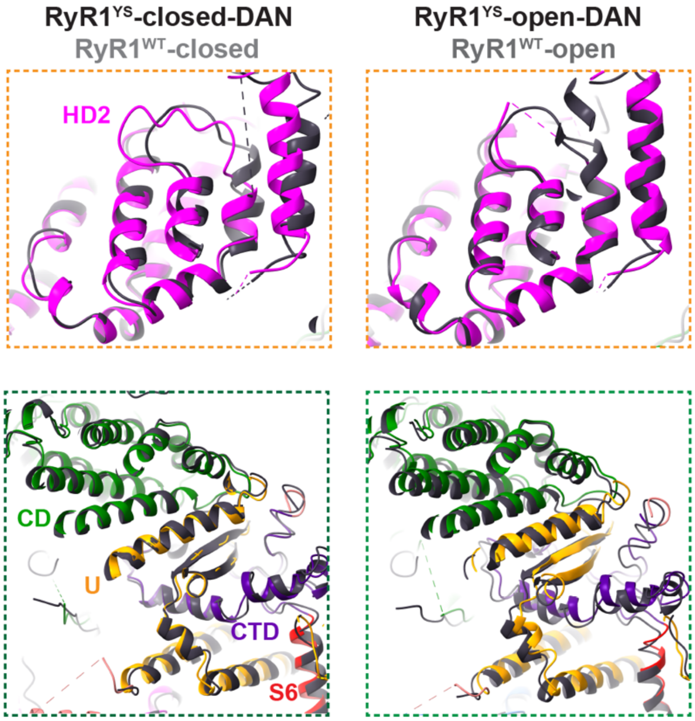

**Figure S10. Dantrolene partially restores the anomalous conformation induced by the YS mutation**

The conformation of mutant RyR1 with dantrolene approached the conformation of WT RyR1 (dark gray PDB IDs: 5TB0 and 5TAL for the closed and open states, respectively) to different extents, depending on the region and the conformation of the channel. This suggests that addition of dantrolene to the mutant set the channel in a distinct state. The regions illustrated by the panels correspond to these shown in Fig. 4; fourfold axis is to the right of the panels.

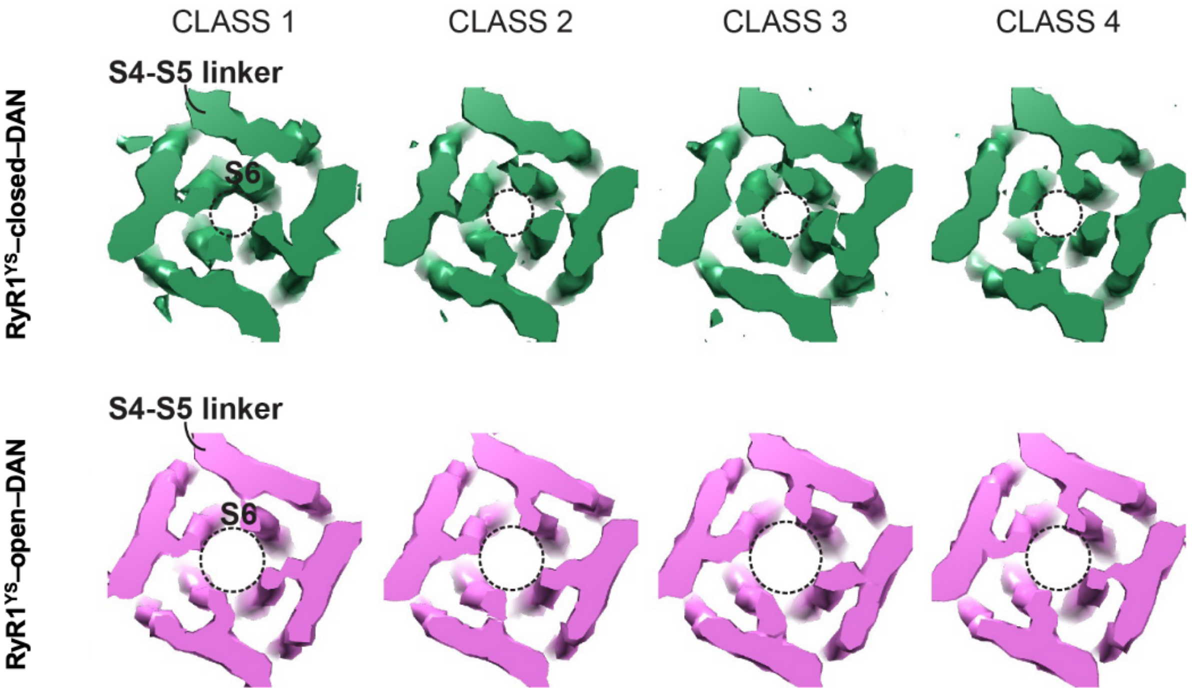

**Figure S11. 3D classification of the two datasets prepared in the presence of EGTA or $Ca^{2+}$**

Slice of the region encompassing the ion gate and S4-S5 linker of resolution/filtered 3D classes for the $RyR1^{YS}$-closed-DAN (green) and $RyR1^{YS}$-open-DAN (pink) datasets indicate uniform populations of closed and open pore, respectively. The structures are low-pass filtered for easier comparison.

**Table S1.** Information about data collection, image processing and model building for the RyR1$^{YS}$-EGTA-DAN and RyR1$^{YS}$- $Ca^{2+}$-DAN datasets.

| | **RyR1$^{YS}$-EGTA-DAN (closed)** | **RyR1$^{YS}$-Ca $^{2+}$-DAN (open)** |
|---|---|---|
| **PURIFICATION** | | |
| Source | HEK293 cells | HEK293 cells |
| Species | Rabbit | Rabbit |
| TmD stabilization | Nanodisc | Nanodisc |
| **STATE** | Closed (2 mM EGTA) | Open (50 $\mu$M $Ca^{2+}$) |
| **BINDING PARTNERS/ LIGANDS** | FKBP12.6, ATP, dantrolene (DAN) | FKBP12.6, ATP, $Ca^{2+}$, dantrolene (DAN) |
| **CRYO-EM** | **EMD-45584 (with refined P1) EMD-45943 (original map)** | **EMD-45585 (with refined P1) EMD-45497 (original map)** |
| Grid type | Quantifoil (Au) | Quantifoil (Au) |
| Sample | 2 μl @ ~2 mg/ml | 2 μl @ ~2 mg/ml |
| Blot time | 1-2 s | 1-2 s |
| Blot force | 1-2 | 1-2 |
| **DATA COLLECTION** | | |
| Pixel size | 1.08 Å | 1.08 Å |
| Magnification | 81,000 | 81,000 |
| Defocus (nominal) | -1.2 to -2.6 μm | -1.2 to -2.6 μm |
| No. of movies collected | 13,992 | 12,212 |
| No. of frames | 40 | 40 |
| Exposure | 2.7 s | 2.7 s |
| Total dose | 50 $e^-/Å^2$ | 50 $e^-/Å^2$ |
| Dose per frame | 1.25 $e^-/Å^2$ | 1.25 $e^-/Å^2$ |
| Camera | K3-Post GIF | K3-Post GIF |
| Energy filter slit width | 10 eV | 10 eV |
| **IMAGE PROCESSING** | | |
| No. of movies after purging | 13,540 | 11,587 |
| No. of images after purging | 12,313 | 11,396 |
| Final no. of particles | 249,034 | 189,847 |
| Resolution | 3.34 Å | 3.23 Å |
| FSC cutoff | 0.143 | 0.143 |
| **MODEL BUILDING** | **PDB: 9CGP** | **PDB: 9CGQ** |
| Ramachandran | | |
| Favored (%) | 90.34 | 91.47 |
| Allowed (%) | 9.63 | 8.50 |
| Outliers (%) | 0.03 | 0.02 |
| Clash score | 9.79 | 8.61 |
